# Early retinal deprivation crossmodally alters nascent subplate circuits and activity in the auditory cortex during the precritical period

**DOI:** 10.1101/2023.02.21.529453

**Authors:** Didhiti Mukherjee, Binghan Xue, Chih-Ting Chen, Minzi Chang, Joseph P. Y. Kao, Patrick O. Kanold

## Abstract

Sensory perturbation in one modality results in adaptive reorganization of neural pathways within the spared modalities, a phenomenon known as “crossmodal plasticity”, which has been examined during or after the classic ‘critical period’. Because peripheral perturbations can alter auditory cortex (ACX) activity and functional connectivity of the ACX subplate neurons (SPNs) even before the classic critical period, called the precritical period, we investigated if retinal deprivation at birth crossmodally alters ACX activity and SPN circuits during the precritical period.

We deprived newborn mice of visual inputs after birth by performing bilateral enucleation. We performed in vivo imaging in the ACX of awake pups during the first two postnatal weeks to investigate cortical activity. We found that enucleation alters spontaneous and sound-evoked activity in the ACX in an age-dependent manner. Next, we performed whole-cell patch clamp recording combined with laser scanning photostimulation in ACX slices to investigate circuit changes in SPNs. We found that enucleation alters the intracortical inhibitory circuits impinging on SPNs shifting the excitation-inhibition balance towards excitation and this shift persists after ear opening. Together, our results indicate that crossmodal functional changes exist in the developing sensory cortices at early ages before the onset of the classic critical period.

## Introduction

*Neural plasticity* allows the brain to adapt to different contexts through rewiring and reorganization. The unpredictable nature of sensory experience prompts different forms of plasticity that enable adaptation to rapidly changing environments (Butz et al., 2009; Hensch & Stryker, 2004; Hubener & Bonhoeffer, 2014; Lorenz, 1935). Although plastic changes in the brain are observed throughout life (Ball & Sekuler, 1982; Sato & Stryker, 2008; Sawtell et al., 2003), they are particularly important during early developmental phases when robust sensory experience from the external environment substantially influences the structural and functional maturation of developing neural structures {for review see (Kolb & Gibb, 2011; Skaliora, 2002)}, thereby making the developing brain extremely vulnerable to loss of environmental stimuli (i.e., sensory deprivation).

Early sensory deprivation in any modality such as visual (Argandona & Lafuente, 1996), auditory (Kral & Eggermont, 2007), or somatosensory (Briner et al., 2010) results in extensive plastic changes within the respective sensory cortices (Argandona & Lafuente, 1996; Briner et al., 2010; Kral & Eggermont, 2007; Kral et al., 2005; Majewska & Sur, 2003; Raggio & Schreiner, 1999), a phenomenon known as “intramodal cortical plasticity”. However, developmental cortical plasticity is not limited to “intramodal” changes. Although the time course for experience-driven sensory development is specific for each modality, sensory perturbations in one modality also result in adaptive reorganization of neural pathways within the spared modalities, including the spared sensory cortices. These adaptive rearrangements within the spared sensory cortices are known as “crossmodal compensatory cortical plasticity” (Bell et al., 2019; Larsen et al., 2009; Mezzera & Lopez-Bendito, 2016; Ramamurthy & Krubitzer, 2018; Striem-Amit et al., 2012; Teichert & Bolz, 2018).

Crossmodal compensatory plasticity within the undeprived sensory cortices, as observed in adults, is most striking when the sensory deprivation is performed very early in development and sustained through the classic “critical period” or is performed during the classic “critical period” [for review see (Bell et al., 2019; Mezzera & Lopez-Bendito, 2016; Teichert & Bolz, 2018)] — a brief developmental time window that commences after thalamic innervation of layer (L) 4 neurons (Barkat et al., 2011; Erzurumlu & Gaspar, 2012) during which the nervous system is robustly shaped by environmental influence (Barkat et al., 2011; Erzurumlu & Gaspar, 2012; Hubel & Wiesel, 1970; Kreile et al., 2011). In altricial experimental animals like rats, cats, mice, ferrets, hamsters etc., eyelids and ear canals open postnatally (Chang & Kanold, 2021; Mezzera & Lopez-Bendito, 2016), and marks the onset of the “classic critical period” (Barkat et al., 2011; Espinosa & Stryker, 2012; Reh et al., 2020). The developmental time before the onset of the classic critical period, frequently termed the “precritical period” has been thought to be governed by genetic factors (Diao et al., 2018; Webber & Raz, 2006) and spontaneous activity (Blankenship & Feller, 2010; Blumberg et al., 2013; Martini et al., 2021; Wang & Bergles, 2015), but devoid of the effects of sensory experience.

Recent evidence, however, demonstrates that sensory deprivation within the same modality (intramodal) can alter cortical circuits and function before the onset of thalamic activation of L4 and the classic critical period (Meng et al., 2021; Mukherjee et al., 2021; Tan et al., 2021). For example, peripheral insult during the precritical period intramodally alters circuit connectivity of the earlier born subplate neurons (SPNs) (Meng et al., 2021; Mukherjee et al., 2021). SPNs are the first targets of thalamocortical inputs before they innervate L4 neurons at the onset of the critical period (Barkat et al., 2011; Friauf et al., 1990; Friauf & Shatz, 1991; Herrmann et al., 1994; Higashi et al., 2002; Molnar et al., 2003; Mukherjee & Kanold, 2022; Zhao et al., 2009). They are essential for the development of thalamocortical and corticocortical connections as well as patterning of the ocular dominance columns and barrels (Ghosh et al., 1990; Ghosh & Shatz, 1992, 1993; Kanold et al., 2003; Kanold & Luhmann, 2010; Kanold & Shatz, 2006; Molnar et al., 2020; Tolner et al., 2012). Peripheral perturbations can alter SPN connections and result in neurodevelopmental disorders (Nagode et al., 2017; Nicolini & Fahnestock, 2018; Sheikh et al., 2019). SPNs in the ACX not only respond to peripheral sound stimulation before L4 neurons are responsive (Wess et al., 2017), but peripheral sound deprivation from birth alters intracortical SPN connectivity in the ACX during the precritical period (Meng et al., 2021; Mukherjee et al., 2021). Moreover, these intramodal changes in SPN circuitry are reflected in altered spontaneous and sound-evoked activity in the infant ACX during this early developmental period. (Meng et al., 2021; Mukherjee et al., 2021).

The developing ACX does not exclusively receive intramodal inputs. Instead, it receives crossmodal inputs from sensory cortices and subcortical structures of other modalities, including structures in the visual pathway (Hanganu-Opatz et al., 2015; Henschke et al., 2018; Kayser & Logothetis, 2007). For example, crossmodal projections from visual thalamus and primary and secondary visual cortices to the ACX are present in neonatal gerbils (Henschke et al., 2018). This raises the questions of whether the developing ACX is vulnerable to a broad range of sensory manipulation, including manipulations of visual function, and if such crossmodal manipulation alters ACX SPN circuits and ACX function before the critical period.

We thus deprived animals of all visual experience and examined the effect on ACX function and ACX SPN circuits. To ensure complete retinal deprivation (spontaneous and light-evoked), we performed bilateral enucleation in newborn mouse pups on postnatal day (P) 1 or 2. To examine ACX function we performed in vivo widefield imaging to record spontaneous and sound-evoked activity in the ACX. To examine ACX SPN circuits we performed whole-cell patch clamp recording from SPNs in thalamocortical slices combined with in vitro laser scanning photostimulation (LSPS). We performed both sets of experiments at P8-9 (before ear opening, precritical period) and P12-15 (around ear opening, onset of critical period). These two ages allowed us to compare the early developmental trajectory of crossmodal changes within the ACX, without retinal input from birth.

We found that complete retinal deprivation at birth results in an increase in spontaneous ACX activity within the first and second postnatal week. Specifically, spontaneous events that are likely of central origin were larger and more frequent after birth enucleation. Similarly, sound-evoked central events were enhanced at both ages, whereas peripheral events remained unchanged. Concurrently, LSPS showed a transient reduction in inhibitory intracortical input to the SPNs at the end of the first postnatal week, which resulted in an imbalance between the excitatory and inhibitory inputs. To our knowledge, this is the first demonstration of functional crossmodal compensatory changes in mammalian sensory cortices before the onset of the classic critical period.

## Materials & Methods

### Animals

All procedures were approved by the Institutional Animal Care and Use Committee (IACUC) at Johns Hopkins University, Baltimore, Maryland, USA. Experiments were performed on C57Bl/6J (JAX strain no. 000664) and Thy1-GCamP6s (JAX strain no. 024275) mouse pups of both sexes aged between P8-P16. Pups were raised with their mothers in standard laboratory cages in a 12-h light/12-h dark condition in the institutional animal colony where lights were turned on at 11:00 am. Food and water were provided *ad libitum*.

### Newborn Bilateral Enucleation

Bilateral enucleation surgeries were performed on C57Bl/6J and Thy1-GCamp6s mouse pups on P1-2 using previously published methods (Deng et al., 2021; Dye et al., 2012). In short, pups were briefly anesthetized with 1-2% inhaled isoflurane (Fluriso, VetOne, Boise, ID). The eyelids were opened with a surgical scalpel blade. The eyeballs were lifted away from the orbit with fine forceps and freed from the optic nerve and surrounding musculature. Following eye removal, eyelids were closed and sealed with surgical glue (Vetbond, 3M, Maplewood, MN). Pups were placed in a plastic box in a warm water bath maintained at 37°C for ~1 h for recovery before returning to their mothers. Sham surgery was performed as control in a cohort of pups at the same age where pups were subjected to anesthesia and revival procedures as described above. All pups were housed with their mothers until they were used for experiments at P8-9 or P12-15 and were weighed routinely to ensure normal thriving.

### In vivo wide field imaging

In vivo wide field imaging was performed following previously published methods (Meng et al., 2021; Mukherjee et al., 2021) on unanesthetized Thy1-GCaMP6s pups at P8-9 (enucleation: *n=9*, sham: *n*=8) and P12-15 (enucleation: *n*=9; sham: *n*=8) after bilateral enucleation or sham surgery on P1-2 (Deng et al., 2021).

#### Surgery

All surgeries were acute, and pups were imaged on the same day. The pup was separated from the litter and was initially anesthetized with 4% inhaled isoflurane (Fluriso, VetOne, Boise, ID). For maintenance, isoflurane concentration was reduced to 2-3%. Throughout surgery, the pup was placed on a heating pad and the body temperature was kept at ~37°C. Depth of anesthesia was monitored every few minutes by observing respiratory pattern and tail-pinch reflex.

The scalp hair was trimmed and the skull overlying the left auditory cortex (ACX) was exposed. Connective tissues were gently removed with the help of a cotton-tipped applicator. Next, a 3D-printed stainless steel headplate (Shapeways, NY) was attached with cyanoacrylate glue on the exposed skull. The headplate was secured to the skull by applying dental cement (C & B Metabond) on the outer perimeter of the headplate. The intact skull was cleaned by topical application of 10% collagenase solution followed by 80% glycerol (Zhao et al., 2018). The pup was lightly wrapped in gauze and placed in a plastic box in a warm water bath maintained at 37°C for ~30 min for recovery.

#### Wide field imaging

After recovery the pup was placed on a far-infrared heating pad (Kent Scientific) over a flat platform, head-fixed and transferred to a sound-proof recording chamber for imaging. In vivo imaging was performed through the intact and cleared skull (Zhao et al., 2018) according to our previously published methods (Francis et al., 2018; Meng et al., 2021; Mukherjee et al., 2021). In brief, blue LED light (470 nm CWL, Thorlabs) was used to excite GCaMP6s fluorescence and emitted light was collected using a tandem lens combination setup. Images were captured with Thorcam software (Thorlabs), which controls a Thorlabs DCC3240M CMOS camera. First, an image of the surface vasculature was captured. Next, the focal plane was advanced to a depth of ~200-400 μm below the surface, where the rest of wide-field imaging was performed across all layers at a rate of 4 frames/sec, with a frame size of 640×512 pixels and a 100-ms exposure time.

Spontaneous activity of the cortex was first recorded for 10 minutes during which no sound was played. Thereafter, we acquired sound-evoked cortical activity. Pure tones of frequencies 4, 8, 16 and 32 kHz were generated using custom software in MATLAB and played from a free field speaker at an intensity of 80 dB sound pressure level (SPL). Each frequency was randomly repeated 12-13 times with an inter-trial interval of 30 s. Each trial consisted of 3-s prestimulus silence, 1-s tone presentation, and 5-s post-stimulus silence.

If the pup exhibited any signs of distress, the experiment was immediately terminated. The pup was euthanized at the end of the experiment.

#### Data analysis

Imaging data was analyzed using custom-written scripts in MATLAB (MathWorks) and as described previously (Meng et al., 2021; Mukherjee et al., 2021). Dimensionality reduction technique was used to perform automatic image segmentation so that pixels with strong temporal correlations across the image were grouped together into single components. We used an autoencoder neural network to perform the dimensionality reduction (Ji Liu et al., 2019). For each region of interest (ROI) we calculated the amplitude and frequency of spontaneous events and the amplitude of events after sound presentation.

For each trial, the response amplitude (ΔF/F) as a function of time for each ROI was defined as ΔF/F = (F–F_0_)/F_0_, where F_0_ corresponds to the baseline fluorescence (defined quantitatively below) and F is the time-varying fluorescence intensity in the ROI. For spontaneous trials F_0_ was calculated by finding the first percentile of fluorescence intensity in sliding windows, the centers of which were equally spaced across the whole trial (window size = 300 frames) and using linear interpolation methods (MATLAB built-in function regress) across all windows. For sound-evoked responses, F_0_ was the first percentile of fluorescence within a 3-s window before tone onset.

For each ROI the averaged ΔF/F was calculated within a 3-s window before and a 3-s window after tone onset for each trial and evaluated with paired-sample t-test comparison between the two averaged ΔF/F across all repeats for each frequency. ROIs that showed a significant increase (p<0.05) in fluorescence after sound presentation at least for one frequency were designated as “responding ROIs”. Responding ROIs are plotted in pseudo-color for ease of visualization.

From the fluorescence traces of each ROI we identified high (H) and low (L)- synchronization events that are distinguished by size in the visual cortex (Siegel et al., 2012) and represent central and peripheral sources respectively (Meng et al., 2021; Siegel et al., 2012). Using the built-in peak detection function (findpeaks) in MATLAB with minimum peak prominence of 0.1 and minimum peak distance of 1 frame we first identified peaks in the fluorescence responses. Next, we used a threshold of 50% ΔF/F to separate L- and H-events. Varying the threshold by ±10% did not affect our results. The response amplitudes of L-/H-events across all the repeats over spontaneous trials or in a period of 3 s after tone onset for each ROI were compared between populations with rank sum test based on Lilliefors test for normality.

### In vitro electrophysiology

In vitro recordings from brain slices were performed as previously described (Cruikshank et al., 2002; Meng et al., 2014; Zhao et al., 2009) on C57Bl/6J mouse pups at P8-9 (enucleation: *n*= 4 pups and 19 cells, sham: *n=* 5 pups and 16 cells) and P12-15 (enucleation: *n=* 5 pups and 15 cells, sham: *n=* 4 pups and 19 cells) after bilateral enucleation or sham surgery was performed on P1-2.

#### Slice preparation

Pups were deeply anesthetized with isofluorane (Fluriso, VetOne, Boise, ID). A block of brain containing primary ACX (A1) and the medial geniculate nucleus (MGN) was isolated and 500 μm thick thalamocortical slices were cut using a vibrating microtome (Leica, Deer Parl, IL) in ice-cold artificial cerebrospinal fluid (ACSF) containing (in mM): 130 NaCl, 3 KCl, 1.25 KH_2_PO_4_, 20 NaHCO_3_, 10 glucose, 1.3 MgSO_4_, 2.5 CaCl2 (pH 7.35-7.4, in 95%O_2_-5%CO_2_). For A1 slices the cutting angle was ~15 degrees from the horizontal plane (lateral raised) as described elsewhere (Cruikshank et al., 2002; Zhao et al., 2009). Thereafter, slices were incubated for ~1 h in ACSF at 30 °C and then were kept at room temperature before they were transferred to the recording chamber.

#### Whole-cell recording

For recording, slices were placed in a chamber on a fixed-stage microscope (Olympus BX51) and superfused at a rate of 2-4 ml/min with high-Mg ACSF (recording solution) at room temperature. High-Mg ACSF reduces spontaneous activity in the slice. The recording solution contained (in mM) 124 NaCl, 5 KCl, 1.23 NaH_2_PO_4_, 26 NaHCO_3_, 10 glucose, 4 MgCl_2_, 4 CaCl2. The location of the recording site in A1 was identified using established landmarks (Cruikshank et al., 2002; Zhao et al., 2009).

Whole-cell recordings were performed with a patch clamp amplifier (Multiclamp 700B, Axon Instruments, San Jose, CA) using pipettes with input resistance of 4-9 MΩ. Data acquisition was performed using National Instruments AD boards and custom software (Ephus) (Suter et al., 2010) written in MATLAB (Mathworks). Voltages were corrected for an estimated junction potential of 10 mV. Electrodes were filled with (in mM) 115 cesium methanesulfonate (CsCH_3_SO_3_), 5 NaF, 10 EGTA, 10 HEPES, 15 CsCl, 3.5 MgATP, 3 QX-314 (pH 7.25, 300 mOsm). Biocytin or Neurobiotin (0.5%) was added to the electrode solution as required. Series resistances were typically 20-25 MΩ.

#### Laser scanning Photostimulation (LSPS)

LSPS was performed using previously published methods (Meng et al., 2015, 2017; Meng et al., 2021; Viswanathan et al., 2017). Briefly, 0.5-1 mM caged glutamate (*N*-(6-nitro-7-coumarylmethyl)-L-glutamate; Ncm-Glu) (Kao, 2006; Muralidharan et al., 2016) was added to the ACSF solution. Without UV light, this compound does not have any effect on neuronal activity (Kao, 2006). UV laser light (500 mW, 355 nm, 1 ms pulses, 100 kHz repetition rate, DPSS lasers) was split by a 33% beam splitter (CVI Melles Griot), attenuated by a Pockels cell (Conoptics), gated with a laser shutter (NM Laser), and coupled into a microscope via scan mirrors (Cambridge Technology) and a dichroic mirror. The laser beam entered the slice axially through the objective (Olympus 10x, 0.3NA/water) and had a diameter of <20 μm. Laser power at the sample was < 25 mW. We typically stimulated up to 30×35 sites spaced 40 μm apart that enabled us to probe areas of 1 mm^2^. Such dense sampling reduced the influence of spontaneous events. Stimuli were applied at 0.5-1 Hz.

#### Data analysis

LSPS was analyzed as described previously (Meng et al., 2015, 2017; Meng et al., 2021; Mukherjee et al., 2021) with custom software written in MATLAB. To detect monosynaptically evoked postsynaltic currents (PSCs), we identified PSCs with onsets in an approximately 50-ms window after the stimulation. This window was chosen as previously observed spiking latency under our recording conditions (Meng et al., 2015, 2017; Meng et al., 2021; Mukherjee et al., 2021).

Recordings were performed at room temperature and in high-Mg^2+^ solution to reduce the probability of multisynaptic inputs. We measured both peak amplitude and transferred charge (integrating the PSC). While the transferred charge may include contributions from multiple events, our prior studies showed a strong correlation between these measures (Meng et al., 2014; Viswanathan et al., 2017). Traces containing a short-latency (< 8 ms) ‘direct’ response were discarded from the analysis as were traces that contained longer latency inward currents of long duration (>50 ms). These currents could sometimes be seen in locations surrounding (<100 μm) areas that gave a ‘direct’ response. Occasionally, some of the ‘direct’ responses contained synaptically evoked responses that we did not separate out, leading to an underestimation of local short-range connections. Cells that did not show any large (>100 pA) direct responses were excluded from the analysis as these could reflect astrocytes or migrating neurons. It is likely that the observed PSCs at each stimulus location represented the activity of multiple presynaptic cells.

Stimulus locations that showed PSCs were deemed connected and we derived binary connection maps. We aligned connection maps for SPNs in the population and averaged connection maps to derive a spatial connection probability map. In these maps the value at each stimulus location indicates the fraction of SPNs that received input from these stimulus locations. Layer boundaries were determined from the infrared images. Next, we derived laminar measures including the input area, integration distance, percentage of excitatory and inhibitory input from each layer, mean and total charge, mean peak and total amplitudes of EPSCs and IPSCs. We calculated the input area for each layer as a measure reflecting the number of presynaptic neurons in each layer projecting to the cell under study. Input area is calculated as the area within each layer that gave rise to PSCs. We also calculated the percentage of input from each layer. Intralaminar integration distance indicates the extent in the rostro-caudal direction that encompasses connected stimulus locations in each layer. Mean charge denotes the average charge of PSCs from each stimulus location in each layer. Peak amplitude measures the mean peak amplitude of EPSCs and IPSCs from each layer. Total charge and amplitude were calculated by multiplying total input area with mean charge and mean peak amplitude, respectively. Since the tonotopic map is largely in the rostro-caudal axis, the intralaminar integration distance reflects integration across the tonotopic axis. While the input area and intralaminar integration are related, the input area shows changes along the columnar (pia-ventricle) axis if more or fewer cells within a tonotopic place are recruited, e.g., only L5 cells vs. L5 and L6 cells. We calculated E/I balance index in each layer for measures of mean charge, mean peak amplitude, total charge and total amplitude as (E-I)/(E+I), thus (AreaE-AreaI)/(Area_E_+Area_I_), resulting in a number that varied between −1 and 1, with 1 indicating dominant excitation and −1 indicating dominant inhibition. Since a measurement of E is not possible close to the soma due to direct responses, we excluded the direct area from both the E and I maps. Thus, this E/I measure does not account for the contribution for cells from close-by locations but does allow analysis of the E/I balance of inputs arising from different layers. We quantified circuit similarity by calculating correlation between connection maps.

#### Statistics

Results are plotted as mean ± SEM unless otherwise stated. Populations from enucleated and sham groups were compared with a rank sum or Mann-Whitney U-test and considered significant if *p* < 0.05.

### Data availability

All data needed to evaluate the conclusions in the paper are presented in the paper and/or the supplemental materials. Additional data related to this paper may be requested from the authors.

## Results

We aimed to investigate crossmodal changes in ACX. The predominant activity in the developing visual system are the spontaneous retinal waves arising from synchronous firing of the retinal ganglion cells (Blankenship & Feller, 2010; Maccione et al., 2014). Importantly, light stimulation through the closed eyelids can alter retinal waves in neonatal mice (Tan et al., 2021; Tiriac et al., 2018). While transgenic mouse lines can alter spontaneous retinal waves at different ages (Bansal et al., 2000), and dark-exposure or lid suture prevents patterned visual inputs from the periphery without affecting retinal spontaneous activity (Chen et al., 2014; Morales et al., 2002), bilateral enucleation irreversibly eliminates both spontaneous and sensory (light-driven) activity at once by instantly and completely removing the eye and retina (Aerts et al., 2014). Although spontaneous thalamic bursts can persist for a certain period after removal of retinal waves (Weliky & Katz, 1999), complete retinal deprivation abolishes early peripheral activity in the visual pathway. Therefore, we chose bilateral enucleation at birth (~P1) to completely remove retinal activity from the earliest ages on.

To examine ACX function after neonatal enucleation, we performed in vivo widefield imaging to record spontaneous and sound-evoked activity in the ACX. Activity in the ACX reflects two sources, peripherally and centrally generated activity. Spontaneous activity originating in the cochlea (Wang & Bergles, 2015) is transmitted via the brainstem and ascending auditory pathways to the developing mouse ACX and is present P7 or earlier (Babola et al., 2018). Centrally generated spontaneous events are also observed in the neonatal ACX (Meng et al., 2021; Mukherjee et al., 2021). Sound-evoked activity originating in the cochlea can be recorded in the ACX by P8 before the onset of the critical period (precritical period, **Fig. 1A**).

**Figure 1.**
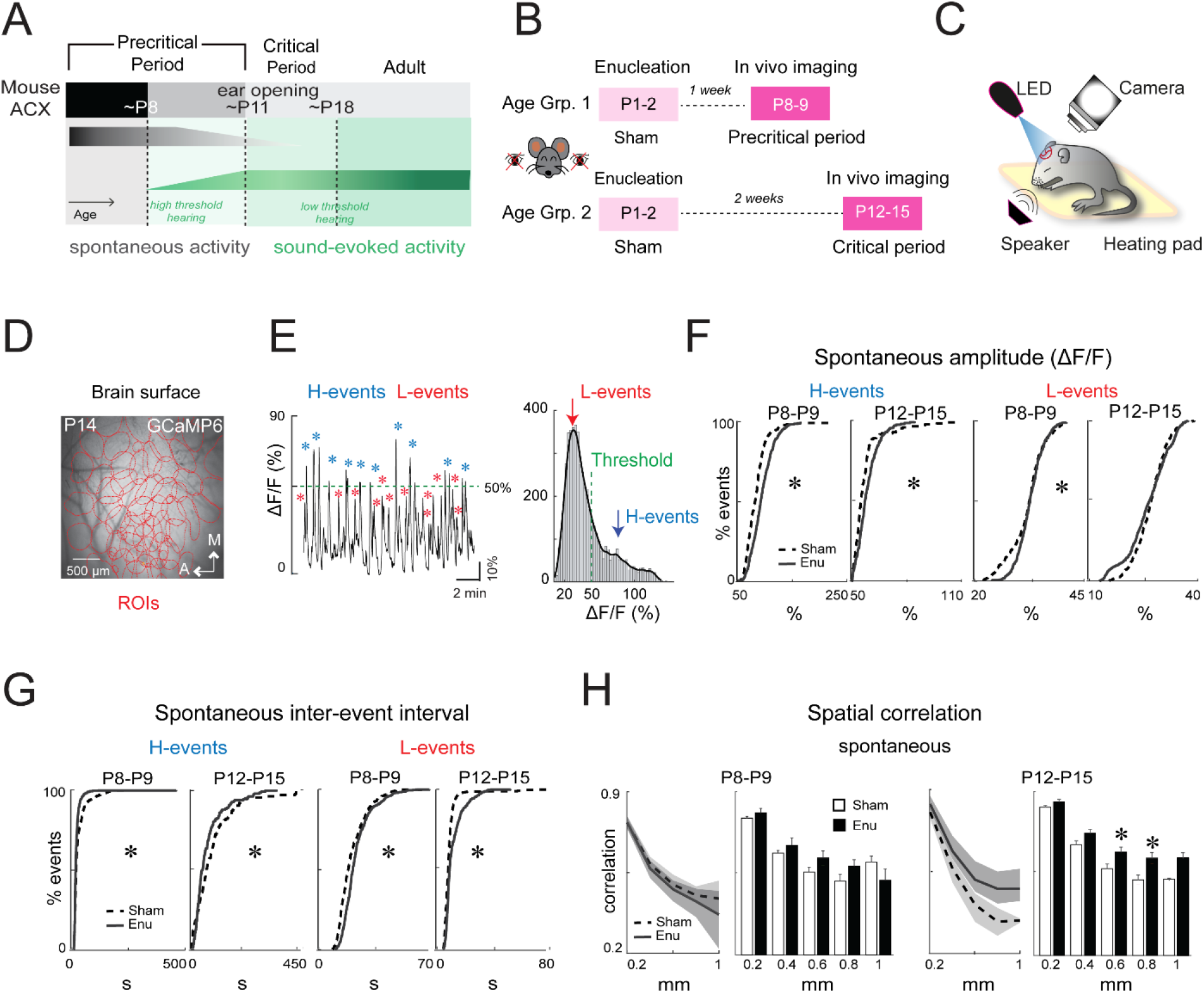
Complete retinal deprivation at birth crossmodally alters spontaneous activity in the ACX at P8-9 and P12-15. **A.** Timeline of ACX development in mice. **B.** Timeline of enucleation surgery and experiments. **C.** Experimental setup for in vivo widefield imaging in awake mouse pups. **D.** Surface of the brain through intact and cleared skull in a representative pup. Open red circles indicate ROIs identified using dimensionality reduction technique. **E.** Raw trace (left) and histogram (right) showing identification of H-and L-events in a representative pup. **F.** Cumulative distribution functions (CDFs) showing peak amplitude of spontaneous H- and L-events in sham (dashed line) and enucleated (solid line) pups at both ages. Amplitude of H-events (left) was higher at both ages (P8-9; p<0.001, Cohen’s d: −0.6, P12-15: p<0.001, Cohen’s d: −0.2) and that of L-events (right) was higher only at P8-9 (p<0.002, Cohen’s d: −0.2) in enucleated pups. **G.** CDFs showing int-erevent interval of H-events (left) was lower (P8-9; p<0.001, Cohen’s d: 0.3, P12-15; p<0.003, Cohen’s d: 0.3) and that of L-events (right) was higher (P8-9; p<0.001, Cohen’s d: −0.3, P12-15: p<0.001, Cohen’s d: −0.4) at both ages in enucleated pups. **H.** Spatial correlation of spontaneous events was higher at 600 μm (p<0.05, Cohen’s d: −0.7) and 800 μm (p<0.05, Cohen’s d: −0.8) distances in enucleated pups at P12-15. ACX: auditory cortex; P: postnatal day; ROI: region of interest; M: medial; A: anterior; H-events: high synchronization events; L-events: low-synchronization events; ΔF/F: change in fluorescence; sham: sham control pups; enu: enucleated pups.

### Altered spontaneous cortical activity at the end of the first and second postnatal weeks after complete retinal deprivation at birth

We performed in vivo widefield imaging in pups expressing calcium indicator GCaMP6 in cortical excitatory neurons under Thy1 promoter (JAX strain no. 024275). We performed imaging before and after the ear canals are open (~P11) to evaluate the impact of low threshold auditory experience (P8-9 enucleation: *n*=9, sham: *n*=8 and P12-15 enucleation: *n*=9; sham: *n*=8, **Fig. 1B, C**). The latter time point coincides with the onset of the critical period (Barkat et al., 2011). In vivo imaging was performed through the intact and cleared skull (Zhao et al., 2018) ~200-400 μm below the brain surface (**Fig. 1D**). We measured both spontaneous and sound evoked activity. Spontaneous activity was imaged for 10 min during which no external sound was played. We then, as in our prior studies, found pixels with strong temporal correlation across the image and grouped them together into single components (ROIs, **Fig. 1D**) (J. Liu et al., 2019; Meng et al., 2021; Mukherjee et al., 2021). As expected, (Meng et al., 2021; Mukherjee et al., 2021), spontaneous activity consisted of high (H) and low (L) synchronization events distinguished by size (**Fig. 1E**). H- and L-events were first demonstrated in the visual cortex and represent central and peripheral sources, respectively (Siegel et al., 2012). Removal of the cochlea reduces the amplitude of ACX L-events consistent with their peripheral origin (Meng et al., 2021; Mukherjee et al., 2021).

Next, we compared the amplitude of spontaneous events in each ROI between the enucleated (enu) and sham control (sham) pups across ages. When compared with the sham controls, the amplitude of the spontaneous H-events was higher in the enucleated pups at P8-9 (medians; sham: 82.3% enu: 92.7%, P<0.001), and at P12-15 (medians; sham: 54.9% enu: 57.1%, P<0.001) (**Fig. 1F**, left), although the effect size was comparatively smaller at the latter age. In contrast, the spontaneous L-events of peripheral origin were mostly unaffected after enucleation. While the amplitude of the spontaneous L-events was marginally higher in enucleated pups at P8-9, the effect size was small (medians; sham: 32.7% enu: 32.9%, P<0.002). The amplitude of spontaneous L-events was similar between groups at P12-15 (medians; sham: 27.1% enu: 27%, P>0.05) (**Fig. 1F**, right). These results suggest that enucleation strengthened spontaneous events of central origin but did not affect spontaneous events originating from the cochlea.

We next compared the frequency of spontaneous H- and L-events (represented as interevent intervals). When compared with sham controls, the frequency of H-events was higher in enucleated pups at both ages (interevent interval medians; P8-9 sham: 19.1 s enu: 16.8 s, P<0.001, P12-15 sham: 64 s enu: 47.3 s, P<0.003) (**Fig. 1G,** left) suggesting an immediate increase in the number of central events after enucleation that persists after the second postnatal week. In contrast, the frequency of L-events was slightly lower in enucleated pups at both ages (interevent interval medians; P8-9 sham: 17.1 s enu: 20.7 s, P<0.001, P12-15 sham: 8 s enu: 9 s P<0.001) (**Fig. 1G**, right), possibly due to a homeostatic compensation. Together these results indicate that early enucleation causes an increase in the amplitude and frequency of centrally generated spontaneous H-events.

Age-related changes in the spontaneous H-and L-event amplitude and inter-event interval have been demonstrated previously. The amplitude of spontaneous H- and L-events are higher at P8-9 than P12-15. The inter-event interval of spontaneous H-events is lower and that of L-events is higher at P8-9 than P12-15 (Meng et al., 2021; Mukherjee et al., 2021). These age-related changes in amplitude and inter-event interval and thus frequency of spontaneous H- and L-events were unaffected after enucleation (**Fig. S1**).

Spontaneous events can correlate distant cortical locations and manipulations of auditory peripheral activity can change these correlations with auditory deprivations resulting in higher correlations of distant ROIs (Meng et al., 2021; Mukherjee et al., 2021). We thus investigated if enucleation altered the spatial correlation of spontaneous activity across ROIs spanning the cortical surface. After enucleation spatial correlations of spontaneous activity was similar between groups at P8-9 (Ps>0.05) (**Fig. 1H, left**), but correlation was higher in enucleated pups at P12-15, specifically at 600 μm (medians: sham-0.49, enu-0.57, P<0.05) and 800 μm (medians: 0.43 sham-0.54, enu-, P<0.05) distances (**Fig. 1H, right**), suggesting distal cortical regions show more correlated activity with age after enucleation.

Together, these results suggest that early retinal deprivation crossmodally alters centrally generated spontaneous activity in the developing ACX during the precritical period. These spontaneous events are more frequent with higher amplitude and involve wider cortical areas with age.

### Transient increase in sound-driven cortical activity at the end of the first postnatal week after complete retinal deprivation at birth

We next examined whether sound-evoked activity in ACX was impacted by enucleation. Sound- evoked responses were recorded after multiple repeats of pure tones of different frequencies were played at 80 dB sound pressure level (Meng et al., 2021; Mukherjee et al., 2021).

Sound-evoked activity was analyzed using previously published methods (Meng et al., 2021; Mukherjee et al., 2021). Sound-responsive ROIs were identified as ROIs that showed significant increase in fluorescence (ΔF/F) after tone onset for at least one frequency (**Fig 2A**) (Meng et al., 2021; Mukherjee et al., 2021). Sound-responsive ROIs were present in both enucleated and sham control pups at both ages. The numbers, total area, and average area of responsive ROIs were similar between groups at both ages (**Fig. S2A**), suggesting cortical area responding to sound stimulation does not differ after enucleation at these ages. There was, however, a significant variability among pups.

**Figure 2.**
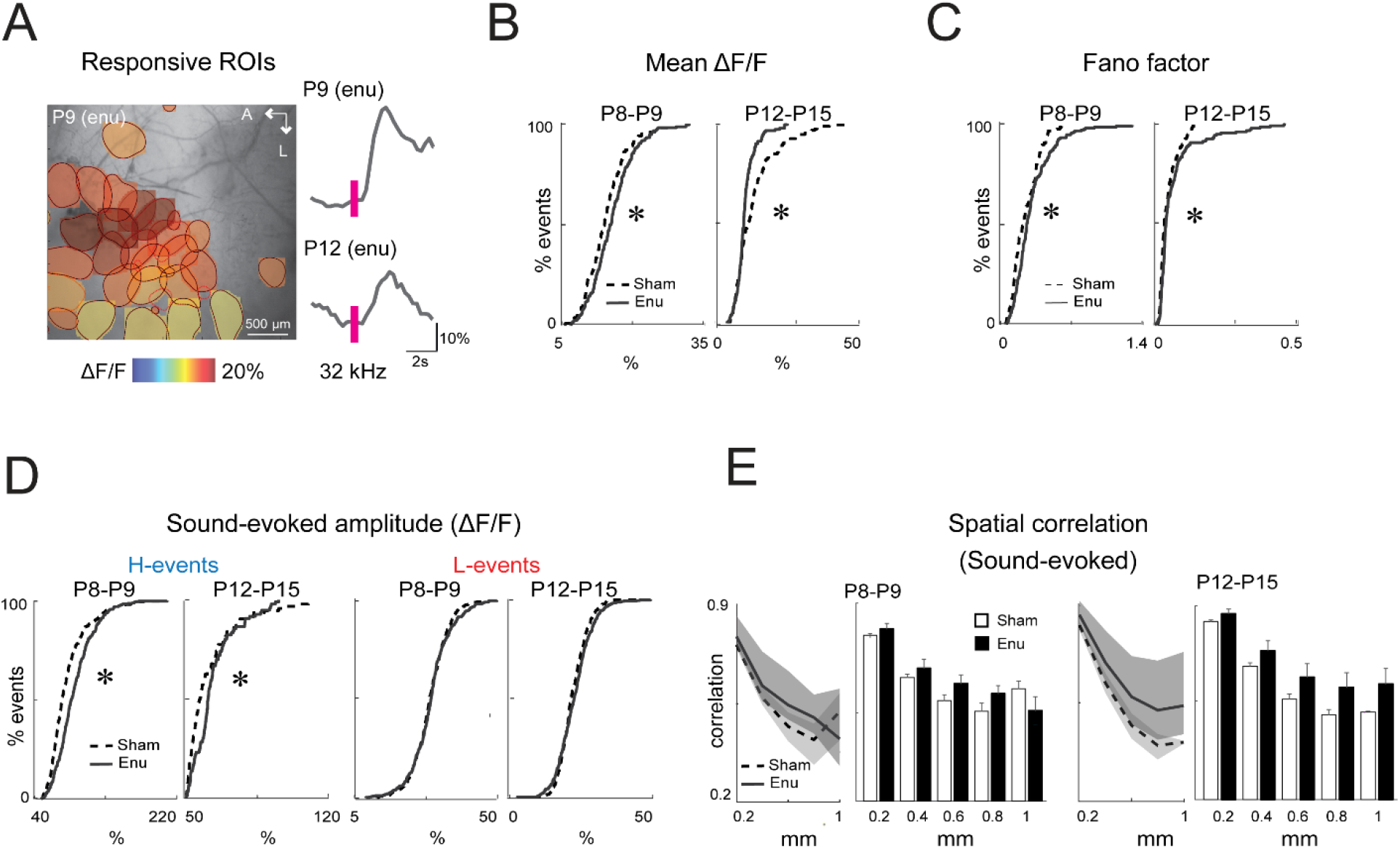
Complete retinal deprivation at birth alters sound-evoked activity in the ACX at P8-9 and P12-15. **A.** Left: filled areas denote responding ROIs that showed significant increase in fluorescence (ΔF/F) within a 3-s window after tone onset. Pseudo-colors indicate mean ΔF/F of responding ROIs. Right: Fluorescence time-course of two representative responding ROIs at P8 (top) and P12 (bottom) after a 32-kHz tone was presented, showing sound-responsiveness in the ACX at these ages. **B.** CDFs showing the mean amplitude of sound-evoked responses was higher (p<0.02, Cohen’s d: −0.1) at P8-9 and lower (p<0.0003, Cohen’s d: 0.6) at P12-15 in enucleated pups. **C.** CDFs showing the fano factor of evoked amplitude was higher (P8-9; p<0.002, Cohen’s d: −0.2, P12-15; p<0.002, Cohen’s d: −0.1) at both ages in enucleated pups suggesting increased variability in the sound-evoked responses, which could be due to an increase in spontaneous activity. **D.** CDFs showing the peak amplitude of sound-evoked H-events (left) was higher at both ages (P8-9; p<0.0001, Cohen’s d: −0.3, P12-15; p<0.004, Cohen’s d: −0.2), whereas that of L- events (right) was not different between groups across ages. **E.** Spatial correlation of sound- evoked events were similar between enucleated and sham control pups across ages.

Next, we calculated the mean fluorescence amplitude within a 3-s window after sound presentation and compared between groups. While the amplitude of the overall sound-evoked responses was marginally higher in the enucleated pups at P8-9 (medians sham: 14.2%; enu: 15.2%, P<0.02) it was marginally lower at P12-15 (medians sham: 11.5%; enu: 10%, P<0.0003) compared their sham controls (**Fig. 2B**). Sound-responsive ROIs can exhibit high variability between trials (Meng et al., 2021; Mukherjee et al., 2021) likely due to weak or immature synapses along the auditory pathway. To identify if enucleation altered this variability, we calculated Fano factor of response amplitude as a measure of variance of sound-evoked activity. Fano factor of response amplitude was slightly higher in enucleated pups at P8-9 (sham: 0.23 enu: 0.25, P<0.002) and at P12-15 (sham: 0.036 enu: 0.043, P<0.002) suggesting enucleation induces some variability in sound responses (**Fig. 2C**), which could be due to an increase in cortically generated spontaneous activity (**Fig. 1**).

Sound-evoked activity comprises both H- and L-events (Meng et al., 2021; Mukherjee et al., 2021). We next compared the amplitude of sound-evoked H- and L-events between groups across ages. The amplitude of sound-evoked H-events was higher in enucleated pups at P8-9 (median: sham: 76.2%; enu: 87.6%, P<0.0001) and at P12-15 (median: sham: 56.8%; enu: 61%, P<0.004) compared to the sham controls. In contrast, the amplitude of the sound-evoked L-events was similar between groups at P8-9 (medians sham: 28.7% enu: 29.1%, P>0.05), and P12-15 (medians sham: 21.7% enu: 22.8%, P>0.05; **Fig. 2D**). These results indicate that early enucleation leads to amplified sound-evoked H-events at both ages without affecting the periphery-driven L-events. Like spontaneous events, the age-related changes in evoked H- and L-event amplitude were unaltered after enucleation (**Fig. S2B**).

We next calculated and compared spatial correlation of sound-evoked activity between groups. The correlation trended higher in enucleated pups at P12-15 but not at P8-9 (**Fig. 2E**).

Together these results suggest that enucleation at birth results in a slight increase in cortical sound-responsiveness involving larger amplitudes. These changes are due to the increase in amplitude of sound-evoked H-events, not affecting periphery originated L-events.

### Inhibitory connections to SPNs are altered at the end of the first postnatal week after enucleation at birth

The observed changes in ACX activity after enucleation might be mirrored by crossmodal changes in functional connectivity in the ACX. Since early born SPNs are the first neurons in ACX to respond to sound they are also vulnerable to a wide range of intramodal sensory and environmental perturbation (Meng et al., 2021; Mukherjee et al., 2021; Sheikh et al., 2019). We thus evaluated the functional connectivity to SPNs after retinal deprivation at birth. To identify circuits to SPNs we used laser-scanning photostimulation (LSPS) combined with whole-cell patch clamp recordings (Meng et al., 2021; Mukherjee et al., 2021; Viswanathan et al., 2017) from SPNs in thalamocortical slices containing ACX (Cruikshank et al., 2002) at P8-9 (enu: *n*= 4 pups and 19 cells, sham: *n=* 5 pups and 16 cells and at P12-15 (enu: *n=* 5 pups and 15 cells, sham: *n=* 4 pups and 19 cells, **Fig. 3A,** left).

**Figure 3.**
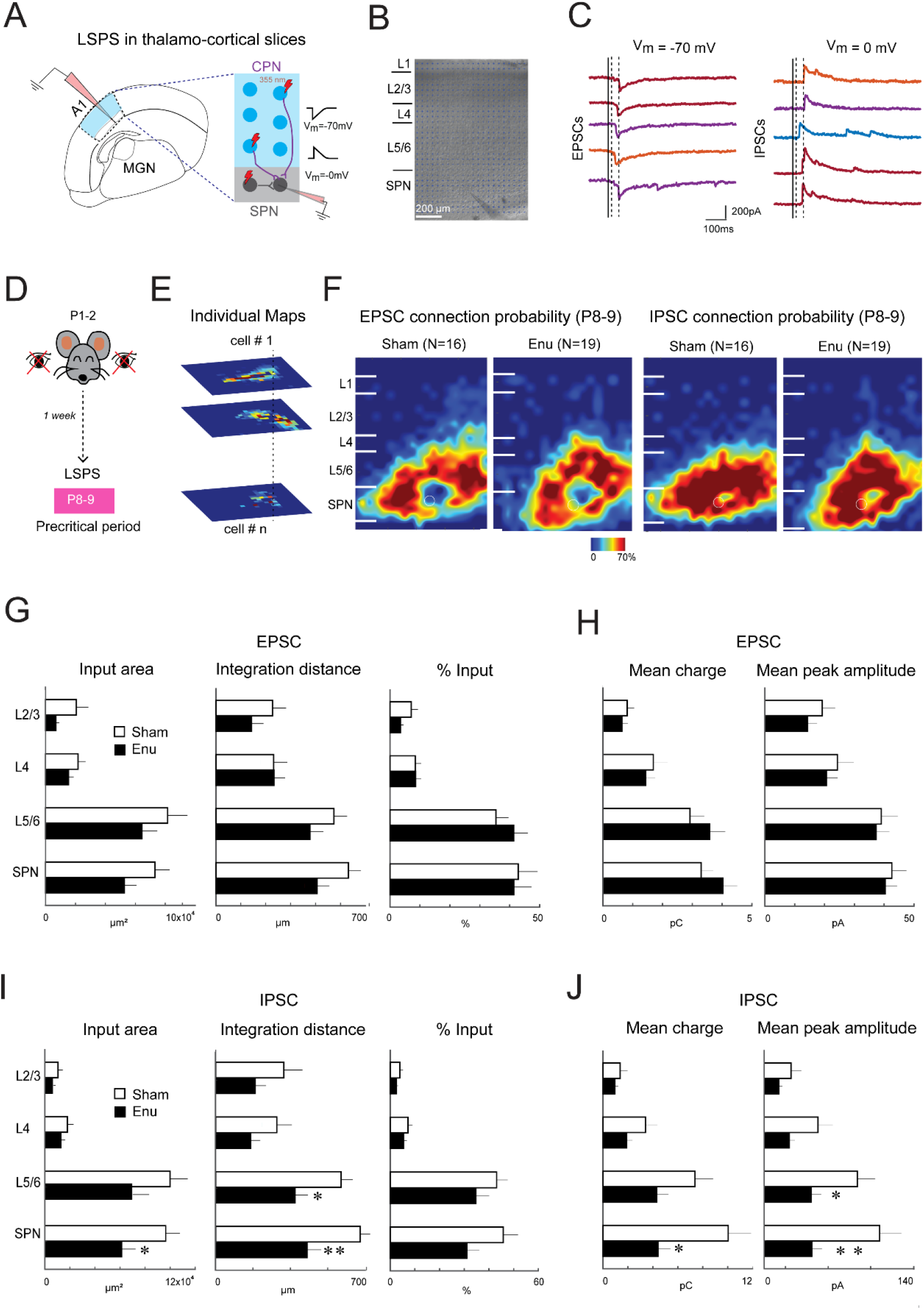
Complete retinal deprivation at birth crossmodally alters inhibitory inputs to SPNs at P8-9. **A.** Left: Cartoon showing whole-cell patch clamp recording from subplate neurons (SPNs) in thalamo-cortical slices. Right: schematic showing laser scanning photostimulation (LSPS). Neurons are activated by laser photolysis (355 nm) of caged glutamate. If stimulated cells are connected to the recorded SPN, evoked excitatory of inhibitory postsynaptic currents (EPSCs and IPSCs) are seen. **B.** Infrared image of an example brain slice with patch pipette on a SPN. **C.** Whole-cell voltage clamp recordings at −70 mV (left) and 0 mV (right) to identify excitatory and inhibitory connections, respectively. Example traces of EPSCs (left) and IPSCs (right) are shown. **D.** Experimental timeline for in vitro experiments during the precritical period after enucleation. **E.** Schematic demonstration of the assembly of connection probability maps by aligning individual maps to the SPN soma (white dashed circle). **F.** Spatial connection probability maps of excitatory and inhibitory connections to the SPN at P8-9. Solid white lines show marginal boundaries. **G.** Bar graphs comparing the layer-specific source area, integration distance and percentage input of excitatory connections between sham (white) and enucleated (black) pups. **H.** Bar graphs showing comparison of layer-specific mean charge and peak amplitude of excitatory connections between sham and enucleated pups. **I.** Bar graphs comparing the layer-specific source area, integration distance and percentage input of inhibitory connections between sham and enucleated pups. Input area from within SPN was lower (p<0.05, Cohen’s d: 0.7) and integration distance from L5/6 and SPN was lower (L5/6; p<0.02, Cohen’s d: 0.8, SPN; p<0.01, Cohen’s d: 1) in enucleated pups. **J.** Bar graphs comparing layer-specific mean charge and peak amplitude of inhibitory connections between sham and enucleated pups. Mean charge from SPNs was lower (p<0.05, Cohen’s d: 0.8) and mean peak amplitude from L5/6 and SP was lower (L5/6; p<0.05, Cohen’s d: 0.8, SPN; p<0.006, Cohen’s d: 1) in enucleated pups. A1: primary auditory cortex; MGN: medial geniculate nucleus; CPN: cortical plate neuron; SPN: subplate neuron; L: layer; Vm: holding voltage.

LSPS measures the functional spatial connection pattern on neurons with ~100 μm resolution over 1 mm^2^ area, which encompasses the whole cortical extent and about 30% of the mouse ACX. LSPS induces action potential in targeted neurons when the laser beam is close to the soma or proximal dendrites. If the activated neuron is connected to the recording neuron, an evoked postsynaptic current (PSC) is observed (**Fig. 3A**, right). For each recorded SPN (**Fig. 3B**), we stimulated 900-1000 locations within the slice and measured the amplitude of evoked excitatory (E) and inhibitory (I) PSCs from each stimulus location by holding the recorded SPN at −70 mV (E_GABA_) or 0 mV (E_Glut_), respectively (**Fig 3C**).

First, we compared spatial connection pattern of excitatory and inhibitory inputs in enucleated pups with their sham counterparts at the end of the first postnatal week (**Fig 3D**). We derived spatial connection maps by plotting the EPSC or IPSC charges at each location. To qualitatively compare spatial connectivity in intra- and interlaminar connections, we aligned the individual connection maps to the position of the patched SPN soma and calculated the percentage of cells receiving inputs from a particular location (**Fig. 3E**). This yields a map of the spatial connection probability of excitatory and inhibitory connections of the patched SPNs. We did not observe any obvious differences in connection probability from these qualitative maps (**Fig. 3F**).

Next, we quantified changes in laminar circuits impinging on the recorded SPNs. For each SPN we calculated the input area, the spatial range of inputs along the tonotopic (rostro-caudal) axis (integration distance) and percentage input received from other cortical layers. The excitatory input, integration distance and percentage input from each layer to the SPNs were similar between both groups (**Fig. 3G**, **S3A,** P>0.05). Mean charges and peak amplitudes of EPSCs were also similar between groups (P>0.05, **Fig. 3H, S3B**).

We next measured the impact of enucleation in inhibitory inputs. Although the percentage of inhibitory inputs from different layers was similar between groups (P>0.05), the input area from within the SPNs was lower (P<0.05) and the integration distance from L5/6 and within the SPNs were also lower (L5/6: P<0.02, SPNs: P<0.01) in enucleated pups (**Fig. 3I, S3C**), indicating a narrower range of intralaminar inputs after enucleation. Mean IPSC charge (P<0.05) from within SPNs and peak IPSC amplitude from L5/6 and within SPNs were lower in enucleated pups (L5/6: P<0.05, SPNs: P<0.006) (**Fig. 3J, S3D)**.

SPNs in ACX show diverse circuit patterns and the diversity of patterns can be sculpted by auditory experience (Meng et al., 2021; Mukherjee et al., 2021). We thus next investigated if enucleation altered SPN diversity by calculating the correlations between connection maps. We found that enucleated pups exhibited lower circuit similarity of excitatory inputs from L5/6 (P<0.0004 **Fig. 4A, S4**). Thus, excitatory inputs from L5/6 to SPNs were more diverse after enucleation. The circuit similarity of inhibitory inputs from L4, L5/6 and SPNs increased after enucleation (L4: P<0.0001, L5/6: P<0.02, SPNs: P<0.0001; **Fig. 4B, S4**). These suggests that early enucleation caused a hypoconnectivity of intralaminar inhibitory cortical connections and increased circuit similarity within the ACX SPNs of at P8-9.

**Figure 4.**
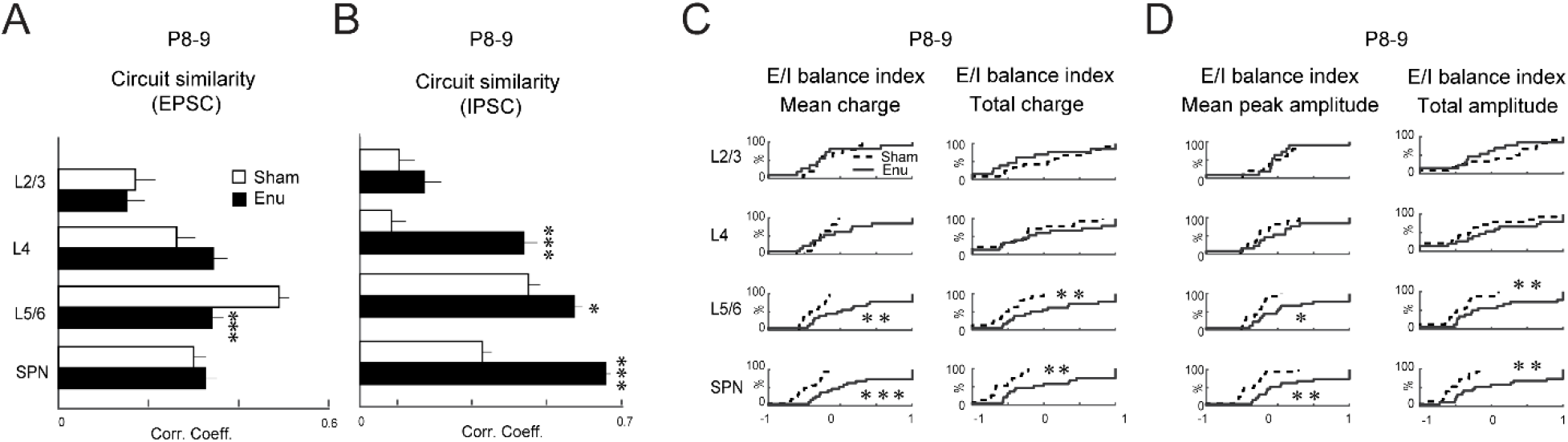
Complete retinal deprivation at birth crossmodally alters inhibitory connection strength to SPNs at P8-9. **A.** Bar graphs showing layer-specific comparison of correlation connection maps of excitatory inputs. Correlation from L5/6 was lower (p<0.0004, Cohen’s d: 0.6) in enucleated pups. **B.** Bar graphs comparing layer-specific correlation connection maps of inhibitory inputs. Correlation from L4, L5/6 and SPNs were higher (L4; p<0.0001, Cohen’s d: −0.8, L5/6; p<0.02, Cohen’s d: −0.5, SPN; p<0.0001, Cohen’s d: −1.4) in enucleated pups. **C.** CDFs showing layer-specific comparison of E/I balance index, [(E-I)/(E+I)], of mean charge and total charge. E/I balance indices of mean and total charge in L5/6 and SPNs were higher (mean charge L5/6; p<0.009, Cohen’s d: −1, mean charge SPN; p<0.001, Cohen’s d: −1, total charge L5/6; p<0.003, Cohen’s d: −1, total charge SPN; p<0.006, Cohen’s d: −1) in enucleated pups. **D.** CDFs show layer-specific comparison of E/I balance index of peak amplitude and total amplitude of postsynaptic currents. E/I balance indices of peak and total amplitude in L5/6 and SPNs were higher (peak amplitude L5/6; p<0.03, Cohen’s d −0.9, peak amplitude SPN: p<0.003, Cohen’s d: −1; total amplitude L5/6: p<0.003, Cohen’s d: −1, total amplitude SPN: p<0.006, Cohen’s d: −1) in enucleated pups.

So far, we investigated the average changes in excitatory and inhibitory connections separately, which could obscure changes in the level of single neurons. We thus quantified excitatory and inhibitory changes at the single cellular level. We calculated E/I balance index, [(E- I)/(E+I)], of mean and peak amplitudes as well as total charges and total amplitudes for each cell to quantify combined excitatory and inhibitory circuit changes. The E/I balance indices based on relative mean and total input charge from L5/6 and SPNs were higher (mean charge: L5/6 P<0.009, SPNs: P<0.001; total charge: L5/6: P<0.003, SPNs: P<0.006) in enucleated pups (**Fig. 4C**). Similarly, E/I balance indices based on peak and total amplitude from L5/6 and SPNs were higher (peak amplitude: L5/6 P<0.03, SPNs: P<0.003; total amplitude: L5/6: P<0.003, SPNs: P<0.006; **Fig. 4D**) in enucleated pups. This indicates a relative change in balance towards excitation after enucleation. Together, these results suggest hypoconnectivity and reduced strengths of inhibitory inputs to ACX SPNs at P8-9 after birth enucleation. These changes are consistent with the overall increase in intracortical activity of excitatory neurons at the end of the first postnatal week as observed in in vivo imaging (**Figs 1 and 2**).

### Excitatory and inhibitory connections to SPNs are altered at the end of the second postnatal week after enucleation at birth

Our functional imaging showed that spontaneous and sound evoked activity in enucleated animals remained altered at P12-P15. We thus next investigated whether SPN circuit changes persisted after ear-opening (P12-15, **Fig. 5A**). The connection probability maps of excitatory inputs at P12-15 showed relatively fewer connections from within the SPNs and more connections from L4 and L2/3 in enucleated pups (**Fig. 5B**). In contrast, the connection probability maps of inhibitory inputs were almost similar in both groups (**Fig. 5C**). We next quantified the laminar circuits impinging on the SPNs. The overall and percentage input area and integration distance of excitatory inputs did not show any difference between groups (P>0.05, **Fig. 5D, S5A**), but the mean charge of excitatory connections from L2/3 was higher (P<0.02) in enucleated pups (**Fig. 5E, S5B**). The laminar measures of inhibitory inputs also did not differ between groups at these ages, except the integration distance from within SPNs were lower (P<0.003) in enucleated pups (**Fig. 5F, S5C**). The mean charge and peak amplitude remained were unaffected (P>0.05, **Fig. 5G, S5D**).

**Figure 5.**
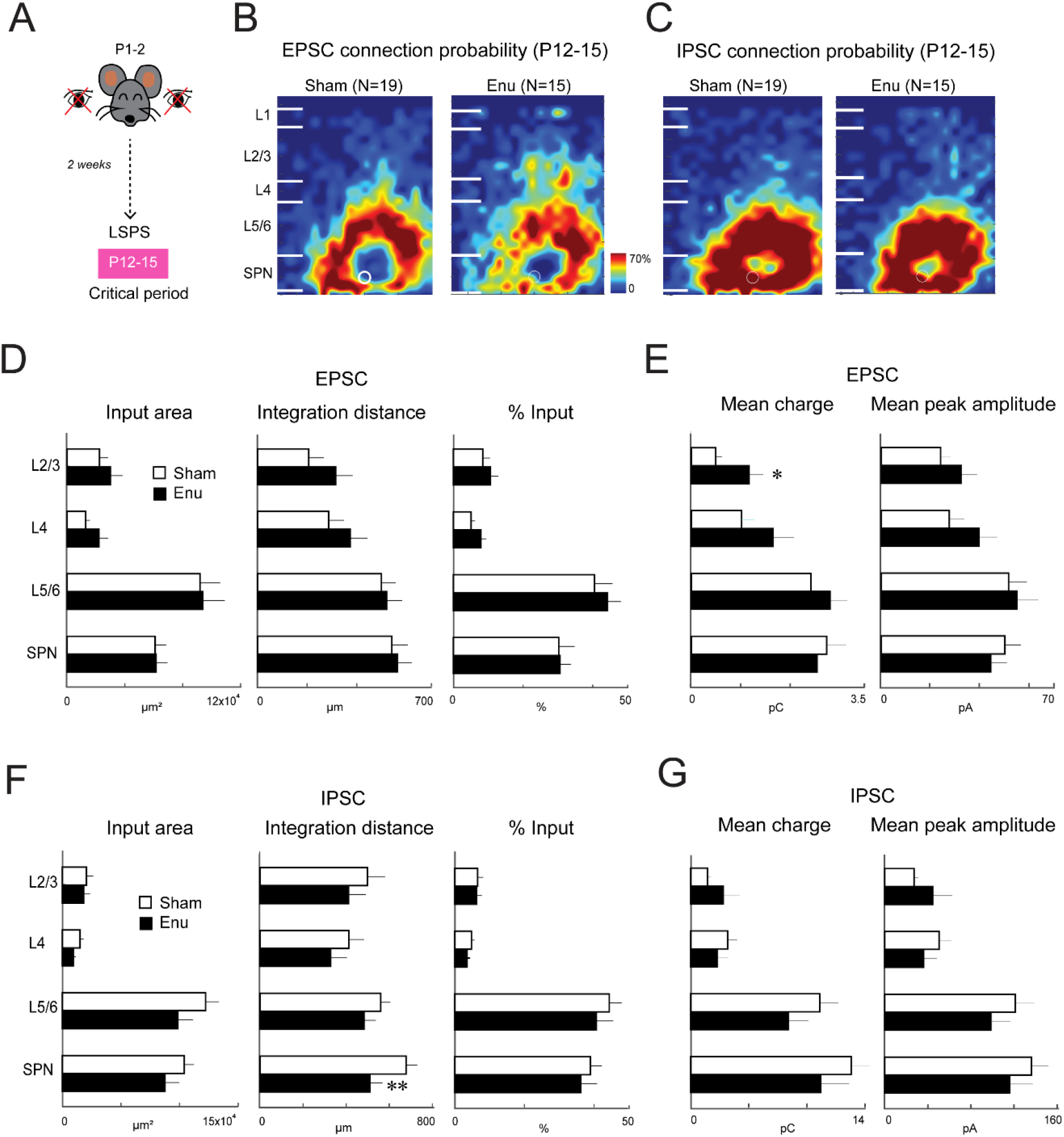
Complete retinal deprivation at birth crossmodally alters excitatory and inhibitory inputs to SPNs at P12-15. **A.** Experimental timeline for in vitro experiments during the critical period after enucleation. **B.** Spatial connection probability map of excitatory connections to the SPN at P12-15. **C.** Spatial connection probability map of inhibitory connections to the SPN at P12-15. **D.** Bar graphs comparing the layer-specific source area, integration distance and percentage input of excitatory connections between sham (white) and enucleated (black) pups. **E.** Bar graphs comparing layer-specific mean charge and peak amplitude of excitatory connections between sham and enucleated pups. Mean charge from L2/3 was higher (p<0.02, Cohen’s d: −0.6) in enucleated pups. **F.** Bar graphs comparing the layer-specific source area, integration distance and percentage input of inhibitory connections between sham and enucleated pups. Integration distance from within SPNs was lower (p<0.003, Cohen’s d: 0.7) in enucleated pups. **G.** Bar graphs comparing layer-specific mean charge and peak amplitude of inhibitory connections between sham and enucleated pups.

We next investigated if enucleation altered SPN diversity at P12-15 by calculating the correlation between connection maps. The circuit similarity of excitatory inputs from L2/3 was higher (P<0.008) and that from within SPNs was lower (P<0.004) in enucleated pups (**Fig. 6A, S6**). This could explain the observed qualitative changes in the maps in **Fig. 5B**. Nonetheless, the circuit similarity of inhibitory inputs from L4 decreased (P<0.008) and that from L5/6 and within SPNs increased (L5/6: P<0.002, SPNs: P<0.0009) after enucleation (**Fig. 6B, S6**).

**Figure 6.**
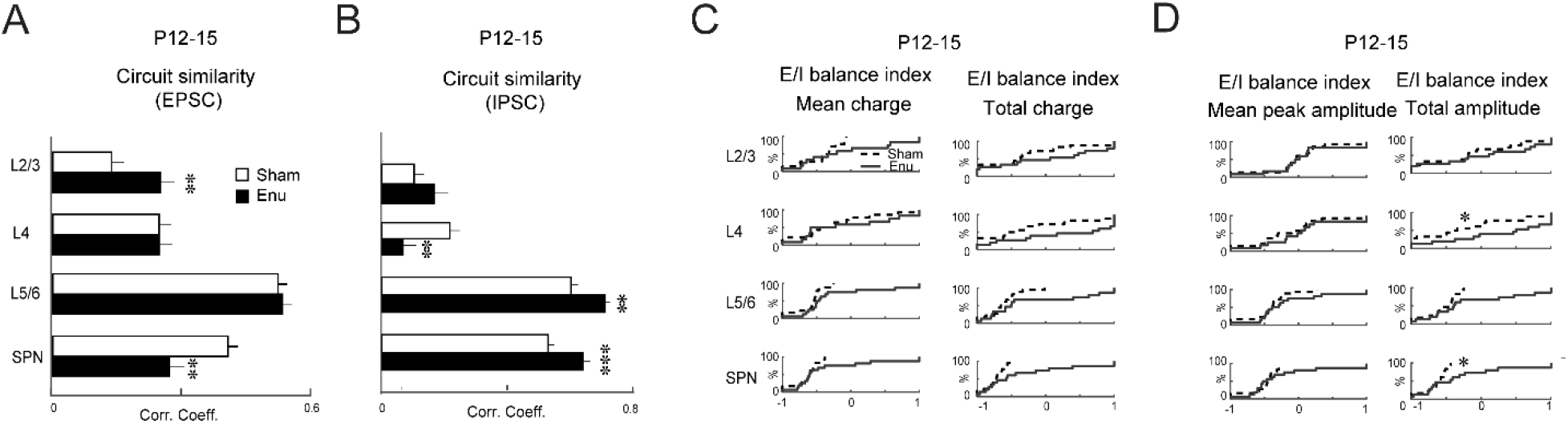
Complete retinal deprivation at birth crossmodally alters excitatory inhibitory connection strength to SPNs at P12-15. **A.** Bar graphs showing layer-specific comparison of correlation connection maps of excitatory inputs. Correlation was higher in L2/3 (p<0.008, Cohen’s d: −0.4) and lower in SPN (p<0.004, Cohen’s d: 0.5) in enucleated pups. **B.** Bar graphs showing layer-specific comparison of correlation connection maps of inhibitory inputs. Correlation was lower in L4 (p<0.008, Cohen’s d: 0.5) and higher in L5/6 and SPN (L5/6; p<0.002, Cohen’s d: −0.5, SPN; p<0.0009, Cohen’s d: −0.5) in enucleated pups. **C.** CDFs show layer-specific comparison of E/I balance index, [(E- I)/(E+I)], of mean charge and total charge. **D.** CDFs show layer-specific comparison of E/I balance index of peak amplitude and total amplitude of postsynaptic currents. E/I balance indices of total amplitude in L4 and SPNs were higher (L4: p<0.05, Cohen’s d: −0.7, SPN; p<0.04, Cohen’s d: −0.7) in enucleated pups.

After ear opening the E/I balance index based on mean and total charges was similar between groups (P>0.05, **Fig. 6C**), whereas that based on total PSC amplitude from L4 and within SPNs was higher (L4: P<0.05, SPNs: P<0.04) in enucleated pups (**Fig. 6D**) suggesting relative increase towards excitation in individual cells.

Together, these results suggest that bilateral enucleation at birth alters excitatory and inhibitory connections to the ACX SPNs at the end of the second postnatal week. Therefore, functional connectivity to ACX SPNs is crossmodally altered within the first two weeks after complete retinal deprivation at birth.

## Discussion

Our results show that complete retinal deprivation from birth crossmodally alters spontaneous and sound-evoked activity as well as functional intracortical connectivity to the SPNs in the developing ACX during the precritical period before the ear canals are open.

Bilateral enucleation at birth crossmodally alters activity and functional connectivity of the ACX in a variety of species (Bell et al., 2019; Dooley & Krubitzer, 2019; Karlen et al., 2006; Mezzera & Lopez-Bendito, 2016; Teichert & Bolz, 2018). However, these crossmodal changes are generally investigated in adults or in developing animals during the “classic critical period”, the brief time window in sensory development imposing heightened sensory influence (Barkat et al., 2011; Erzurumlu & Gaspar, 2012; Hubel & Wiesel, 1970; Kreile et al., 2011). Here we show that early retinal deprivation crossmodally alters activity and functional connectivity in the developing mouse ACX within the first postnatal week, even before the onset of the classic critical period (~P11 in rats, ~P12 in mice) (Barkat et al., 2011; de Villers-Sidani et al., 2007). Such immediate and short-term crossmodal cortical changes have been reported in the developing rodent somatosensory system using gene expression and anatomical techniques. For example, enucleation at birth in mice results in positional shift in developmentally regulated gene (e.g., *ephrin A5*) expression and inter-neocortical projections at the somatosensory-visual border in only 10 days (Dye et al., 2012). In rats enucleated at birth, a crossmodal expansion of the barrel fields is also observed within the somatosensory cortex at the end of the first postnatal week (Fetter-Pruneda et al., 2013) — days before the onset of active whisking (Grant et al., 2012). To our knowledge, our results are the first demonstration of experience-dependent early crossmodal functional changes within the auditory system, days before the ear canals are open. These changes may serve as a substrate for homeostatic sensory compensation in early development.

Our results show that enucleated animals show reduced inhibitory connections in the first postnatal weak causing an imbalance between excitation and inhibition towards excitation, in particular for inputs from deep layers and that this imbalance is still present after ear opening. The observed imbalance towards excitation is consistent with our observation of increased spontaneous activity and increase spatial spread of activity correlations.

We observe a transient interlaminar circuit changes to SPNs. The overall inhibitory inputs from L5/6 and within the SPN were significantly compromised in enucleated pups at P8-9, which led to an imbalance between the excitatory and inhibitory inputs and a resultant increase in excitation. Such crossmodal changes in intralaminar cortical circuits has also been demonstrated in older mice after dark exposure (Meng et al., 2015, 2017). Visual deprivation has been shown to weaken intramodal synaptic inhibition within the visual cortex (Gabbott & Stewart, 1987; Morales et al., 2002). We here show that crossmodal manipulations also have the ability to change inhibitory circuits in the developing ACX.

We find that circuits from deep L5/6 neurons and from within SP are most impacted by retinal deprivation. Thus, we speculate that retinal input is relayed to ACX via these circuits. These crossmodal inputs to ACX could be intracortical and/or carried by thalamic afferents including afferents from visual thalamus, which innervates the developing ACX (Henschke et al., 2018).

Early crossmodal changes in the developing ACX included a persistent increase in amplitude and frequency of spontaneous events likely of cortical origin during the first and second postnatal weeks. Such instantaneous increase in spontaneous activity is also observed in thalamo-recipient L4 neurons in A1 after a brief visual deprivation (dark exposure for 7 days) in adult mice (Petrus et al., 2014). Additionally, dark exposure for a short period of time significantly enhances spontaneous raw local field potential (LFP) oscillations as well as β oscillations in A1 L4 neurons in juvenile (P28) rats, suggesting crossmodal increase in excitability of A1 neurons and higher synchronization of activity within a widespread cortical area (Pan et al., 2018). Immediate changes in spontaneous activity is also observed within days of bilateral enucleation in the primary somatosensory cortex of newborn rats (Martinez-Mendez et al., 2019), which may mechanistically underlie the crossmodal barrel expansion as reported elsewhere (Fetter-Pruneda et al., 2013). Therefore, the observed increase in frequency and amplitude of spontaneous H-events could possibly underlie the crossmodal expansion of the ACX as observed after bilateral enucleation (Kahn & Krubitzer, 2002). Additionally, increased spontaneous activity could enhance the excitability of the cortical neurons to underlie the enhanced crossmodal sound activation and perception later in life, as observed in adult animals after visual deprivation (Korte & Rauschecker, 1993; Petrus et al., 2014; Rauschecker & Kniepert, 1994) and in congenitally blind individuals [for review see (Bell et al., 2019)].

In addition to intracortical changes, alteration in thalamo-cortical activity could also contribute to the observed changes. Crossmodal thalamo-cortical changes are observed after visual deprivation in the auditory pathway at older ages. For example, an increase in LFP oscillation power in the auditory thalamus (medial geniculate nucleus) is observed in juvenile (P28) rats after bilateral enucleation (Pan et al., 2018). Dark exposure results in strengthening of thalamocortical synapses to A1 L4 neurons in adult mice (Petrus et al., 2014). However, we observe only small changes in L-events suggesting that changes in additory thalamocortical circuits after enucleation are likely to be very small. In addition to changes in ascending auditory pathway, the small transient changes in ACX L-events could also result from changes within the visual thalamus (lateral geniculate nucleus, LGN) that sends afferent inputs to the developing ACX (Henschke et al., 2018). LGN is activated after visual deprivation (Bhandari et al., 2022; Giasafaki et al., 2022), and experiences rewiring of cortico-thalamic projections after enucleation (Frangeul et al., 2016; Giasafaki et al., 2022; Grant et al., 2016). Together, we reason that the functional changes we observe are mostly due to changes in intracortical circuits.

Our results demonstrate that in addition to intramodal rewiring of SPN circuits after sensory deprivation (Meng et al., 2021; Mukherjee et al., 2021), compensatory crossmodal rearrangement of intra- and inter-laminar circuits are observed in the cortical SPNs before the onset of the classic critical period. This suggests that the earlier-born SPNs are the earliest cortical substrate for a wide range of early experience-dependent plasticity.

Identifying the developmental emergence of crossmodal changes and their underlying mechanisms is critically important to implement effective therapeutic measures to recover or restore early loss of sensory function in children. For example, in children with congenital or other forms of deafness, the earlier the cochlear implant is implemented, the better the chances of restoring hearing before it is taken over by other modalities (Hoff et al., 2019; Peixoto et al., 2013). Given the key role of SPNs in shaping the development of cortical layer 4 and beyond (Kanold et al., 2003; Kanold & Shatz, 2006), early transient crossmodal influences onto SPNs might have the potential to affect later development of other cortical areas even beyond the initial crossmodal effect. Thus, limiting or exploiting such early crossmodal interactions might be beneficial for therapeutic interventions.

## Notes

**Funding**, This work was supported by NIH R01DC009607 (POK) and NIH R01GM056481 (JPYK).

**Competing interests**, The authors report no competing interests.

### Competing Interest Statement

The authors have declared no competing interest.

## References

Aerts, J., Nys, J., &Arckens, L. (2014). A highly reproducible and straightforward method to perform in vivo ocular enucleation in the mouse after eye opening. J Vis Exp(92), e51936. https://doi.org/10.3791/51936

Argandona, E. G., & Lafuente, J. V. (1996). Effects of dark-rearing on the vascularization of the developmental rat visual cortex. Brain Res, 732(1-2), 43–51.https://doi.org/10.1016/0006-8993(96)00485-4

Babola, T. A., Li, S., Gribizis, A., Lee, B. J., Issa, J. B., Wang, H. C., Crair, M. C., & Bergles, D. E. (2018). Homeostatic Control of Spontaneous Activity in the Developing Auditory System. Neuron, 99(3), 511–524 e515. https://doi.org/10.1016/j.neuron.2018.07.004

Ball, K., & Sekuler, R. (1982). A specific and enduring improvement in visual motion discrimination. Science, 218(4573), 697–698. https://doi.org/10.1126/science.7134968

Bansal, A., Singer, J. H., Hwang, B. J., Xu, W., Beaudet, A., & Feller, M. B. (2000). Mice lacking specific nicotinic acetylcholine receptor subunits exhibit dramatically altered spontaneous activity patterns and reveal a limited role for retinal waves in forming ON and OFF circuits in the inner retina. J Neurosci, 20(20), 7672–7681. https://www.ncbi.nlm.nih.gov/pubmed/11027228

Barkat, T. R., Polley, D. B., & Hensch, T. K. (2011). A critical period for auditory thalamocortical connectivity. Nat Neurosci, 14(9), 1189–1194. https://doi.org/10.1038/nn.2882

Bell, L., Wagels, L., Neuschaefer-Rube, C., Fels, J., Gur, R. E., & Konrad, K. (2019). The Cross-Modal Effects of Sensory Deprivation on Spatial and Temporal Processes in Vision and Audition: A Systematic Review on Behavioral and Neuroimaging Research since 2000. Neural Plast, 2019, 9603469. https://doi.org/10.1155/2019/9603469

Bhandari, A., Ward, T. W., Smith, J., & Van Hook, M. J. (2022). Structural and Functional Plasticity in the Dorsolateral Geniculate Nucleus of Mice following Bilateral Enucleation. Neuroscience, 488, 44–59. https://doi.org/10.1016/j.neuroscience.2022.01.029

Blankenship, A. G., & Feller, M. B. (2010). Mechanisms underlying spontaneous patterned activity in developing neural circuits. Nat Rev Neurosci, 11(1), 18–29. https://doi.org/10.1038/nrn2759

Blumberg, M. S., Marques, H. G., & Iida, F. (2013). Twitching in sensorimotor development from sleeping rats to robots. Curr Biol, 23(12), R532–537. https://doi.org/10.1016/j.cub.2013.04.075

Briner, A., De Roo, M., Dayer, A., Muller, D., Kiss, J. Z., & Vutskits, L. (2010). Bilateral whisker trimming during early postnatal life impairs dendritic spine development in the mouse somatosensory barrel cortex. J Comp Neurol, 518(10), 1711–1723. https://doi.org/10.1002/cne.22297

Butz, M., Worgotter, F., & van Ooyen, A. (2009). Activity-dependent structural plasticity. Brain Res Rev, 60(2), 287–305. https://doi.org/10.1016/j.brainresrev.2008.12.023

Chang, M., & Kanold, P. O. (2021). Development of Auditory Cortex Circuits. J Assoc Res Otolaryngol, 22(3), 237–259. https://doi.org/10.1007/s10162-021-00794-3

Chen, X. J., Rasch, M. J., Chen, G., Ye, C. Q., Wu, S., & Zhang, X. H. (2014). Binocular input coincidence mediates critical period plasticity in the mouse primary visual cortex. J Neurosci, 34(8), 2940–2955. https://doi.org/10.1523/JNEUROSCI.2640-13.2014

Cruikshank, S. J., Rose, H. J., & Metherate, R. (2002). Auditory thalamocortical synaptic transmission in vitro. J Neurophysiol, 87(1), 361–384. http://www.ncbi.nlm.nih.gov/entrez/query.fcgi?cmd=Retrieve&db=PubMed&dopt=Citation&listuids=11784756

de Villers-Sidani, E., Chang, E. F., Bao, S., & Merzenich, M. M. (2007). Critical period window for spectral tuning defined in the primary auditory cortex (A1) in the rat. J Neurosci, 27(1), 180–189. https://doi.org/10.1523/JNEUROSCI.3227-06.2007

Deng, R., Kao, J. P. Y., & Kanold, P. O. (2021). Aberrant development of excitatory circuits to inhibitory neurons in the primary visual cortex after neonatal binocular enucleation. Sci Rep, 11(1), 3163. https://doi.org/10.1038/s41598-021-82679-2

Diao, Y., Chen, Y., Zhang, P., Cui, L., & Zhang, J. (2018). Molecular guidance cues in the development of visual pathway. Protein Cell, 9(11), 909–929. https://doi.org/10.1007/s13238-017-0490-7

Dooley, J. C., & Krubitzer, L. A. (2019). Alterations in cortical and thalamic connections of somatosensory cortex following early loss of vision. J Comp Neurol, 527(10), 1675–1688.

https://doi.org/10.1002/cne.24582

Dye, C. A., Abbott, C. W., & Huffman, K. J. (2012). Bilateral enucleation alters gene expression and intraneocortical connections in the mouse. Neural Dev, 7, 5. https://doi.org/10.1186/1749-8104-7-5

Erzurumlu, R. S., & Gaspar, P. (2012). Development and critical period plasticity of the barrel cortex. Eur J Neurosci, 35(10), 1540–1553. https://doi.org/10.1111/j.1460-9568.2012.08075.

x Espinosa, J. S., & Stryker, M. P. (2012). Development and plasticity of the primary visual cortex. Neuron, 75(2), 230–249. https://doi.org/10.1016/j.neuron.2012.06.009

Fetter-Pruneda, I., Geovannini-Acuna, H., Santiago, C., Ibarraran-Viniegra, A. S., Martinez-Martinez, E., Sandoval-Velasco, M., Uribe-Figueroa, L., Padilla-Cortes, P., Mercado-Celis, G., & Gutierrez-Ospina, G. (2013). Shifts in developmental timing, and not increased levels of experience-dependent neuronal activity, promote barrel expansion in the primary somatosensory cortex of rats enucleated at birth. PLoS One, 8(1), e54940. https://doi.org/10.1371/journal.pone.0054940

Francis, N. A., Winkowski, D. E., Sheikhattar, A., Armengol, K., Babadi, B., & Kanold, P. O. (2018). Small Networks Encode Decision-Making in Primary Auditory Cortex. Neuron, 97(4), 885–897 e886. https://doi.org/10.1016/j.neuron.2018.01.019

Frangeul, L., Pouchelon, G., Telley, L., Lefort, S., Luscher, C., & Jabaudon, D. (2016). A cross-modal genetic framework for the development and plasticity of sensory pathways. Nature, 538(7623), 96–98. https://doi.org/10.1038/nature19770

Friauf, E., McConnell, S. K., & Shatz, C. J. (1990). Functional synaptic circuits in the subplate during fetal and early postnatal development of cat visual cortex. J Neurosci, 10(8), 2601–2613.

Friauf, E., & Shatz, C. J. (1991). Changing patterns of synaptic input to subplate and cortical plate during development of visual cortex. J Neurophysiol, 66(6), 2059–2071.

Gabbott, P. L., & Stewart, M. G. (1987). Quantitative morphological effects of dark-rearing and light exposure on the synaptic connectivity of layer 4 in the rat visual cortex (area 17). Exp Brain Res, 68(1), 103–114. https://doi.org/10.1007/BF00255237

Ghosh, A., Antonini, A., McConnell, S. K., & Shatz, C. J. (1990). Requirement for subplate neurons in the formation of thalamocortical connections. Nature, 347(6289), 179–181.

Ghosh, A., & Shatz, C. J. (1992). Involvement of subplate neurons in the formation of ocular dominance columns. Science, 255(5050), 1441–1443.

Ghosh, A., & Shatz, C. J. (1993). A role for subplate neurons in the patterning of connections from thalamus to neocortex. Development, 117(3), 1031–1047. http://www.ncbi.nlm.nih.gov/cgi-bin/Entrez/referer? http://www.cob.org.uk/Development/117/03/dev9138.html

Giasafaki, C., Grant, E., Hoerder-Suabedissen, A., Hayashi, S., Lee, S., & Molnar, Z. (2022). Cross-hierarchical plasticity of corticofugal projections to dLGN after neonatal monocular enucleation. J Comp Neurol, 530(7), 978–997. https://doi.org/10.1002/cne.25304

Grant, E., Hoerder-Suabedissen, A., & Molnar, Z. (2016). The Regulation of Corticofugal Fiber Targeting by Retinal Inputs. Cereb Cortex, 26(3), 1336–1348. https://doi.org/10.1093/cercor/bhv315

Grant, R. A., Mitchinson, B., & Prescott, T. J. (2012). The development of whisker control in rats in relation to locomotion. Dev Psychobiol, 54(2), 151–168. https://doi.org/10.1002/dev.20591

Hanganu-Opatz, I. L., Rowland, B. A., Bieler, M., & Sieben, K. (2015). Unraveling Cross-Modal Development in Animals: Neural Substrate, Functional Coding and Behavioral Readout. Multisens Res, 28(1-2), 33–69. https://doi.org/10.1163/22134808-00002477

Hensch, T. K., & Stryker, M. P. (2004). Columnar architecture sculpted by GABA circuits in developing cat visual cortex. Science, 303(5664), 1678–1681. http://www.ncbi.nlm.nih.gov/entrez/query.fcgi?cmd=Retrieve&db=PubMed&dopt=Citation&listuids=15017001

Henschke, J. U., Oelschlegel, A. M., Angenstein, F., Ohl, F. W., Goldschmidt, J., Kanold, P. O., & Budinger, E. (2018). Early sensory experience influences the development of multisensory thalamocortical and intracortical connections of primary sensory cortices. Brain Struct Funct, 223(3), 1165–1190. https://doi.org/10.1007/s00429-017-1549-1

Herrmann, K., Antonini, A., & Shatz, C. J. (1994). Ultrastructural evidence for synaptic interactions between thalamocortical axons and subplate neurons. Eur J Neurosci, 6(11), 1729–1742. https://doi.org/10.1111/j.1460-9568.1994.tb00565.x

Higashi, S., Molnar, Z., Kurotani, T., & Toyama, K. (2002). Prenatal development of neural excitation in rat thalamocortical projections studied by optical recording. Neuroscience, 115(4), 1231–1246. https://doi.org/10.1016/s0306-4522(02)00418-9

Hoff, S., Ryan, M., Thomas, D., Tournis, E., Kenny, H., Hajduk, J., & Young, N. M. (2019). Safety and Effectiveness of Cochlear Implantation of Young Children, Including Those With Complicating Conditions. Otol Neurotol, 40(4), 454–463. https://doi.org/10.1097/MAO.0000000000002156

Hubel, D. H., & Wiesel, T. N. (1970). The period of susceptibility to the physiological effects of unilateral eye closure in kittens. J Physiol, 206(2), 419–436. https://doi.org/10.1113/jphysiol.1970.sp009022

Hubener, M., & Bonhoeffer, T. (2014). Neuronal plasticity: beyond the critical period. Cell, 159(4), 727–737. https://doi.org/10.1016/j.cell.2014.10.035

Kahn, D. M., & Krubitzer, L. (2002). Massive cross-modal cortical plasticity and the emergence of a new cortical area in developmentally blind mammals. Proc Natl Acad Sci U S A, 99(17), 11429–11434. https://doi.org/10.1073/pnas.162342799

Kanold, P. O., Kara, P., Reid, R. C., & Shatz, C. J. (2003). Role of subplate neurons in functional maturation of visual cortical columns. Science, 301(5632), 521–525. https://doi.org/10.1126/science.1084152

Kanold, P. O., & Luhmann, H. J. (2010). The subplate and early cortical circuits. Annu Rev Neurosci, 33, 23–48. https://doi.org/10.1146/annurev-neuro-060909-153244

Kanold, P. O., & Shatz, C. J. (2006). Subplate neurons regulate maturation of cortical inhibition and outcome of ocular dominance plasticity. Neuron, 51(5), 627–638. https://doi.org/10.1016/j.neuron.2006.07.008

Kao, J. P. Y. (2006). Caged molecules: principles and practical considerations. In C. Gerfen, A. Holmes, M. Rogawski, D. Sibley, P. Skolnick, & S. Wray (Eds.), Current protocols in neuroscience (Vol. Unit 6.20). Wiley.

Karlen, S. J., Kahn, D. M., & Krubitzer, L. (2006). Early blindness results in abnormal corticocortical and thalamocortical connections. Neuroscience, 142(3), 843–858. https://doi.org/10.1016/j.neuroscience.2006.06.055

Kayser, C., & Logothetis, N. K. (2007). Do early sensory cortices integrate cross-modal information? Brain Struct Funct, 212(2), 121–132. https://doi.org/10.1007/s00429-007-0154-0

Kolb, B., & Gibb, R. (2011). Brain plasticity and behaviour in the developing brain. J Can Acad Child Adolesc Psychiatry, 20(4), 265–276. https://www.ncbi.nlm.nih.gov/pubmed/22114608

Korte, M., & Rauschecker, J. P. (1993). Auditory spatial tuning of cortical neurons is sharpened in cats with early blindness. J Neurophysiol, 70(4), 1717–1721. https://doi.org/10.1152/jn.1993.70.4.1717

Kral, A., & Eggermont, J. J. (2007). What’s to lose and what’s to learn: development under auditory deprivation, cochlear implants and limits of cortical plasticity. Brain Res Rev, 56(1), 259–269. https://doi.org/10.1016/j.brainresrev.2007.07.021

Kral, A., Tillein, J., Heid, S., Hartmann, R., & Klinke, R. (2005). Postnatal cortical development in congenital auditory deprivation. Cereb Cortex, 15(5), 552–562. https://doi.org/10.1093/cercor/bhh156

Kreile, A. K., Bonhoeffer, T., & Hubener, M. (2011). Altered visual experience induces instructive changes of orientation preference in mouse visual cortex. J Neurosci, 31(39), 13911–13920. https://doi.org/10.1523/JNEUROSCI.2143-11.2011

Larsen, D. D., Luu, J. D., Burns, M. E., & Krubitzer, L. (2009). What are the Effects of Severe Visual Impairment on the Cortical Organization and Connectivity of Primary Visual Cortex? Front Neuroanat, 3, 30. https://doi.org/10.3389/neuro.05.030.2009

Liu, J., Whiteway, M. R., Sheikhattar, A., Butts, D. A., Babadi, B., & Kanold, P. O. (2019). Parallel processing of sound dynamics across mouse auditory cortex via spatially patterned thalamic inputs and distinct areal intracortical circuits. Cell reports, 27(3), 872–885. e877.

Liu, J., Whiteway, M. R., Sheikhattar, A., Butts, D. A., Babadi, B., & Kanold, P. O. (2019). Parallel Processing of Sound Dynamics across Mouse Auditory Cortex via Spatially Patterned Thalamic Inputs and Distinct Areal Intracortical Circuits. Cell Rep, 27(3), 872-885 e877. https://doi.org/10.1016/j.celrep.2019.03.069

Lorenz, K. (1935). Der Kumpan in der Umwelt des Vogels. Journal für Ornithologie, 83(3), 289–413. https://doi.org/10.1007/BF01905572

Maccione, A., Hennig, M. H., Gandolfo, M., Muthmann, O., van Coppenhagen, J., Eglen, S. J., Berdondini, L., & Sernagor, E. (2014). Following the ontogeny of retinal waves: pan-retinal recordings of population dynamics in the neonatal mouse. J Physiol, 592(7), 1545–1563. https://doi.org/10.1113/jphysiol.2013.262840

Majewska, A., & Sur, M. (2003). Motility of dendritic spines in visual cortex in vivo: changes during the critical period and effects of visual deprivation. Proc Natl Acad Sci U S A, 100(26), 16024–16029. https://doi.org/10.1073/pnas.2636949100

Martinez-Mendez, R., Perez-Torres, D., Gomez-Chavarin, M., Padilla-Cortes, P., Fiordelisio, T., & Gutierrez-Ospina, G. (2019). Bilateral enucleation at birth modifies calcium spike amplitude, but not frequency, in neurons of the somatosensory thalamus and cortex: Implications for developmental cross-modal plasticity. IBRO Rep, 7, 108–116. https://doi.org/10.1016/j.ibror.2019.11.003

Martini, F. J., Guillamon-Vivancos, T., Moreno-Juan, V., Valdeolmillos, M., & Lopez-Bendito, G. (2021). Spontaneous activity in developing thalamic and cortical sensory networks. Neuron, 109(16), 2519–2534. https://doi.org/10.1016/j.neuron.2021.06.026

Meng, X., Kao, J. P., & Kanold, P. O. (2014). Differential signaling to subplate neurons by spatially specific silent synapses in developing auditory cortex. J Neurosci, 34(26), 8855–8864. https://doi.org/10.1523/JNEUROSCI.0233-14.2014

Meng, X., Kao, J. P., Lee, H. K., & Kanold, P. O. (2015). Visual Deprivation Causes Refinement of Intracortical Circuits in the Auditory Cortex. Cell Rep, 12(6), 955–964. https://doi.org/10.1016/j.celrep.2015.07.018

Meng, X., Kao, J. P., Lee, H. K., & Kanold, P. O. (2017). Intracortical Circuits in Thalamorecipient Layers of Auditory Cortex Refine after Visual Deprivation. eNeuro, 4(2). https://doi.org/10.1523/ENEURO.0092-17.2017

Meng, X., Mukherjee, D., Kao, J. P. Y., & Kanold, P. O. (2021). Early peripheral activity alters nascent subplate circuits in the auditory cortex. Sci Adv, 7(7). https://doi.org/10.1126/sciadv.abc9155

Mezzera, C., & Lopez-Bendito, G. (2016). Cross-modal plasticity in sensory deprived animal models: From the thalamocortical development point of view. J Chem Neuroanat, 75(Pt A), 32–40. https://doi.org/10.1016/j.jchemneu.2015.09.005

Molnar, Z., Kurotani, T., Higashi, S., Yamamoto, N., & Toyama, K. (2003). Development of functional thalamocortical synapses studied with current source-density analysis in whole forebrain slices in the rat. Brain Res Bull, 60(4), 355–371. http://www.ncbi.nlm.nih.gov/entrez/query.fcgi?cmd=Retrieve&db=PubMed&dopt=Citation&listuids=12781324

Molnar, Z., Luhmann, H. J., & Kanold, P. O. (2020). Transient cortical circuits match spontaneous and sensory-driven activity during development. Science, 370(6514). https://doi.org/10.1126/science.abb2153

Morales, B., Choi, S. Y., & Kirkwood, A. (2002). Dark rearing alters the development of GABAergic transmission in visual cortex. J Neurosci, 22(18), 8084–8090. https://www.ncbi.nlm.nih.gov/pubmed/12223562

Mukherjee, D., & Kanold, P. O. (2022). Changing subplate circuits: Early activity dependent circuit plasticity. Front Cell Neurosci, 16, 1067365. https://doi.org/10.3389/fncel.2022.1067365

Mukherjee, D., Meng, X., Kao, J. P. Y., & Kanold, P. O. (2021). Impaired Hearing and Altered Subplate Circuits During the First and Second Postnatal Weeks of Otoferlin-Deficient Mice. Cereb Cortex. https://doi.org/10.1093/cercor/bhab383

Muralidharan, S., Dirda, N. D., Katz, E. J., Tang, C. M., Bandyopadhyay, S., Kanold, P. O., & Kao, J. P. (2016). Ncm, a Photolabile Group for Preparation of Caged Molecules: Synthesis and Biological Application. PLoS One, 11(10), e0163937. https://doi.org/10.1371/journal.pone.0163937

Nagode, D. A., Meng, X., Winkowski, D. E., Smith, E., Khan-Tareen, H., Kareddy, V., Kao, J. P. Y., & Kanold, P. O. (2017). Abnormal Development of the Earliest Cortical Circuits in a Mouse Model of Autism Spectrum Disorder. Cell Rep, 18(5), 1100–1108. https://doi.org/10.1016/j.celrep.2017.01.006

Nicolini, C., & Fahnestock, M. (2018). The valproic acid-induced rodent model of autism. Exp Neurol, 299(Pt A), 217–227. https://doi.org/10.1016/j.expneurol.2017.04.017

Pan, P., Zhou, Y., Fang, F., Zhang, G., & Ji, Y. (2018). Visual deprivation modifies oscillatory activity in visual and auditory centers. Anim Cells Syst (Seoul), 22(3), 149–156. https://doi.org/10.1080/19768354.2018.1474801

Peixoto, M. C., Spratley, J., Oliveira, G., Martins, J., Bastos, J., & Ribeiro, C. (2013). Effectiveness of cochlear implants in children: long term results. Int J Pediatr Otorhinolaryngol, 77(4), 462–468. https://doi.org/10.1016/j.ijporl.2012.12.005

Petrus, E., Isaiah, A., Jones, A. P., Li, D., Wang, H., Lee, H. K., & Kanold, P. O. (2014). Crossmodal induction of thalamocortical potentiation leads to enhanced information processing in the auditory cortex. Neuron, 81(3), 664–673. https://doi.org/10.1016/j.neuron.2013.11.023

Raggio, M. W., & Schreiner, C. E. (1999). Neuronal responses in cat primary auditory cortex to electrical cochlear stimulation. III. Activation patterns in short- and long-term deafness. J Neurophysiol, 82(6), 3506–3526. https://doi.org/10.1152/jn.1999.82.6.3506

Ramamurthy, D. L., & Krubitzer, L. A. (2018). Neural Coding of Whisker-Mediated Touch in Primary Somatosensory Cortex Is Altered Following Early Blindness. J Neurosci, 38(27), 6172–6189. https://doi.org/10.1523/JNEUROSCI.0066-18.2018

Rauschecker, J. P., & Kniepert, U. (1994). Auditory localization behaviour in visually deprived cats. Eur J Neurosci, 6(1), 149–160. https://doi.org/10.1111/j.1460-9568.1994.tb00256.x

Reh, R. K., Dias, B. G., Nelson, C. A., 3rd, Kaufer, D., Werker, J. F., Kolb, B., Levine, J. D., & Hensch, T. K. (2020). Critical period regulation across multiple timescales. Proc Natl Acad Sci U S A, 117(38), 23242–23251. https://doi.org/10.1073/pnas.1820836117

Sato, M., & Stryker, M. P. (2008). Distinctive features of adult ocular dominance plasticity. J Neurosci, 28(41), 10278–10286. https://doi.org/10.1523/JNEUROSCI.2451-08.2008

Sawtell, N. B., Frenkel, M. Y., Philpot, B. D., Nakazawa, K., Tonegawa, S., & Bear, M. F. (2003). NMDA receptor-dependent ocular dominance plasticity in adult visual cortex. Neuron, 38(6), 977–985. https://doi.org/10.1016/s0896-6273(03)00323-4

Sheikh, A., Meng, X., Liu, J., Mikhailova, A., Kao, J. P. Y., McQuillen, P. S., & Kanold, P. O. (2019). Neonatal Hypoxia-Ischemia Causes Functional Circuit Changes in Subplate Neurons. Cereb Cortex, 29(2), 765–776. https://doi.org/10.1093/cercor/bhx358

Siegel, F., Heimel, J. A., Peters, J., & Lohmann, C. (2012). Peripheral and central inputs shape network dynamics in the developing visual cortex in vivo. Curr Biol, 22(3), 253–258. https://doi.org/10.1016/j.cub.2011.12.026

Skaliora, I. (2002). Experience-dependent plasticity in the developing brain. International Congress Series, 1241, 313–320. https://doi.org/10.1016/S0531-5131(02)00616-7

Striem-Amit, E., Bubic, A., & Amedi, A. (2012). Neurophysiological Mechanisms Underlying Plastic Changes and Rehabilitation following Sensory Loss in Blindness and Deafness. In M. M. Murray & M. T. Wallace (Eds.), The Neural Bases of Multisensory Processes. https://www.ncbi.nlm.nih.gov/pubmed/22593863

Suter, B. A., O’Connor, T., Iyer, V., Petreanu, L. T., Hooks, B. M., Kiritani, T., Svoboda, K., & Shepherd, G. M. (2010). Ephus: multipurpose data acquisition software for neuroscience experiments. Front Neural Circuits, 4, 100. https://doi.org/10.3389/fncir.2010.00100

Tan, L., Ringach, D. L., Zipursky, S. L., & Trachtenberg, J. T. (2021). Vision is required for the formation of binocular neurons prior to the classical critical period. Curr Biol. https://doi.org/10.1016/j.cub.2021.07.053

Teichert, M., & Bolz, J. (2018). How Senses Work Together: Cross-Modal Interactions between Primary Sensory Cortices. Neural Plast, 2018, 5380921. https://doi.org/10.1155/2018/5380921

Tiriac, A., Smith, B. E., & Feller, M. B. (2018). Light Prior to Eye Opening Promotes Retinal Waves and Eye-Specific Segregation. Neuron, 100(5), 1059-1065 e1054. https://doi.org/10.1016/j.neuron.2018.10.011

Tolner, E. A., Sheikh, A., Yukin, A. Y., Kaila, K., & Kanold, P. O. (2012). Subplate neurons promote spindle bursts and thalamocortical patterning in the neonatal rat somatosensory cortex. J Neurosci, 32(2), 692–702. https://doi.org/10.1523/JNEUROSCI.1538-11.2012

Viswanathan, S., Sheikh, A., Looger, L. L., & Kanold, P. O. (2017). Molecularly Defined Subplate Neurons Project Both to Thalamocortical Recipient Layers and Thalamus. Cereb Cortex, 27(10), 4759–4768. https://doi.org/10.1093/cercor/bhw271

Wang, H. C., & Bergles, D. E. (2015). Spontaneous activity in the developing auditory system. Cell Tissue Res, 361(1), 65–75. https://doi.org/10.1007/s00441-014-2007-5

Webber, A., & Raz, Y. (2006). Axon guidance cues in auditory development. Anat Rec A Discov Mol Cell Evol Biol, 288(4), 390–396. https://doi.org/10.1002/ar.a.20299

Weliky, M., & Katz, L. C. (1999). Correlational structure of spontaneous neuronal activity in the developing lateral geniculate nucleus in vivo. Science, 285(5427), 599–604. https://doi.org/10.1126/science.285.5427.599

Wess, J. M., Isaiah, A., Watkins, P. V., & Kanold, P. O. (2017). Subplate neurons are the first cortical neurons to respond to sensory stimuli. Proc Natl Acad Sci U S A, 114(47), 12602–12607. https://doi.org/10.1073/pnas.1710793114

Zhao, C., Kao, J. P., & Kanold, P. O. (2009). Functional excitatory microcircuits in neonatal cortex connect thalamus and layer 4. J Neurosci, 29(49), 15479–15488. https://doi.org/10.1523/JNEUROSCI.4471-09.2009

Zhao, Y. J., Yu, T. T., Zhang, C., Li, Z., Luo, Q. M., Xu, T. H., & Zhu, D. (2018). Skull optical clearing window for in vivo imaging of the mouse cortex at synaptic resolution. Light Sci Appl, 7, 17153. https://doi.org/10.1038/lsa.2017.153

