## Supplemental Figures 1-6 for "Early retinal deprivation crossmodally alters nascent subplate circuits and activity in the auditory cortex during the precritical period"

#### Supplemental figures and legends

Figures S1-S6

Figure S1

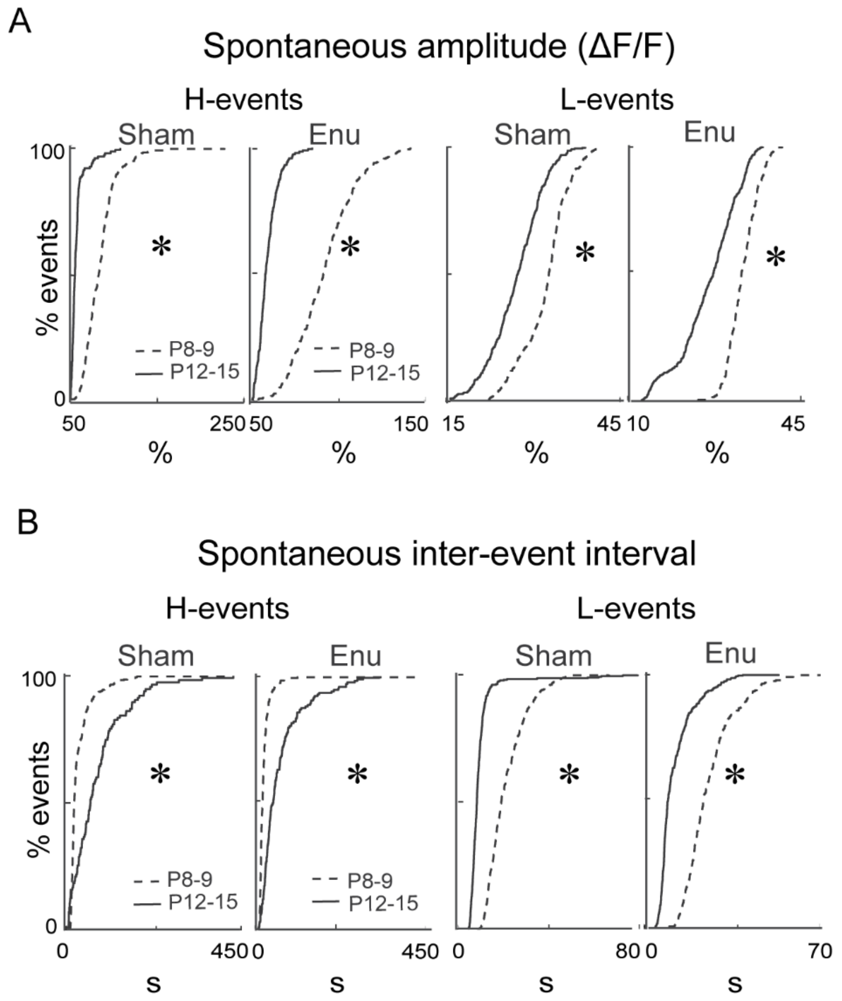


**Figure S1. Age-related changes in spontaneous H- and L-events are not altered after enucleation.**

**A.** CDFs showing the amplitude of H- (left) and L-events (right) are lower at P12-15 than P8-9 in sham controls and enucleated pups. **B.** CDFs showing the inter-event interval of H-events (left) are higher and that of L-events (right) are lower at P12-15 than P8-9 in sham controls and enucleated pups.

Figure S2

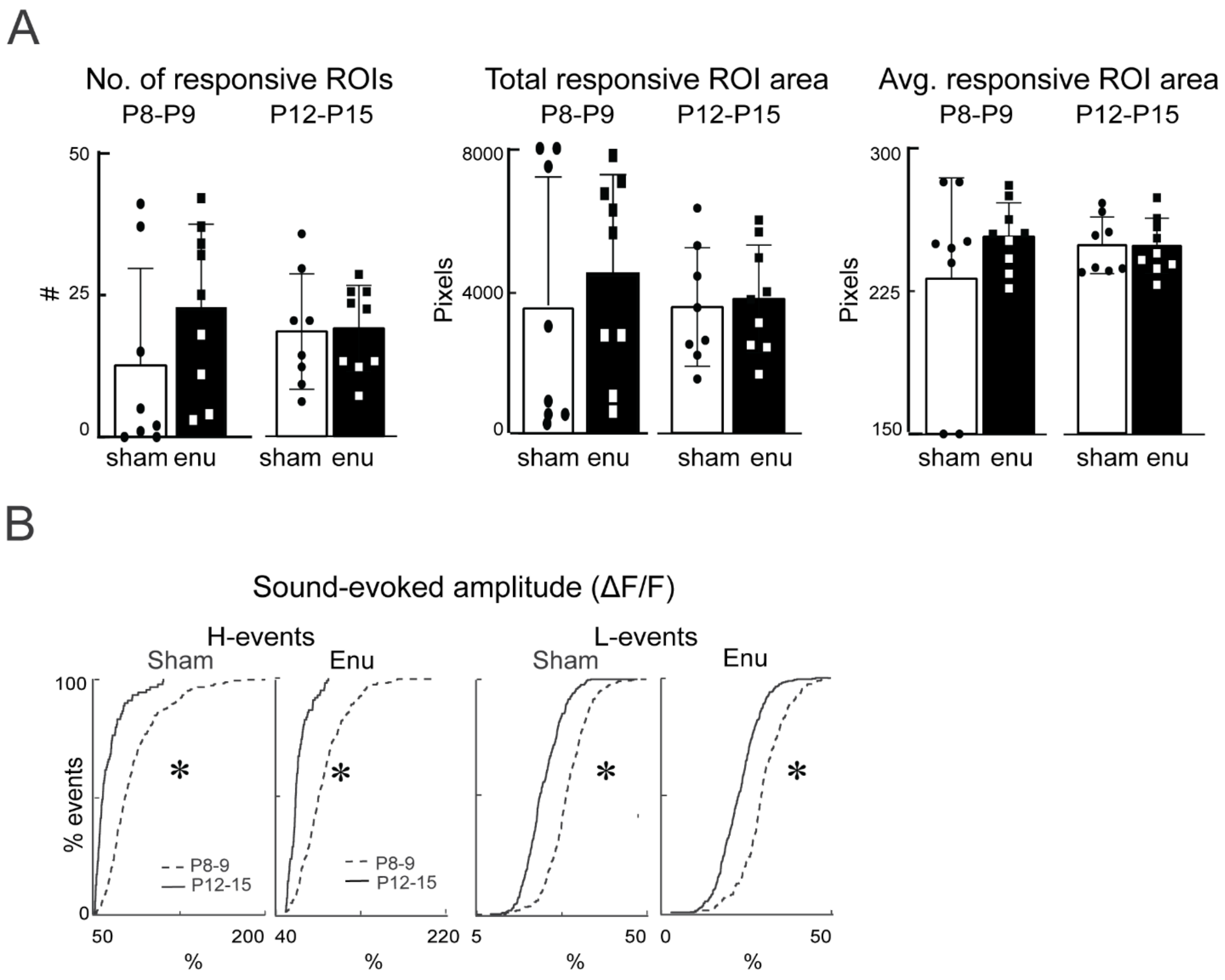


**Figure S2. Sound-responsive ROIs and age-related changes in sound-evoked H- and L-events are not altered after enucleation.**

**A.** Bar graphs showing the number (left), total responding ROI area (middle), and average responding ROI area (right) did not differ between groups across ages. Subjective variability was observed. **B.** CDFs showing the amplitude of sound-evoked H- (left) and L-events (right) are lower at P12-15 than P8-9 in sham controls and enucleated pups.

### Figure S3

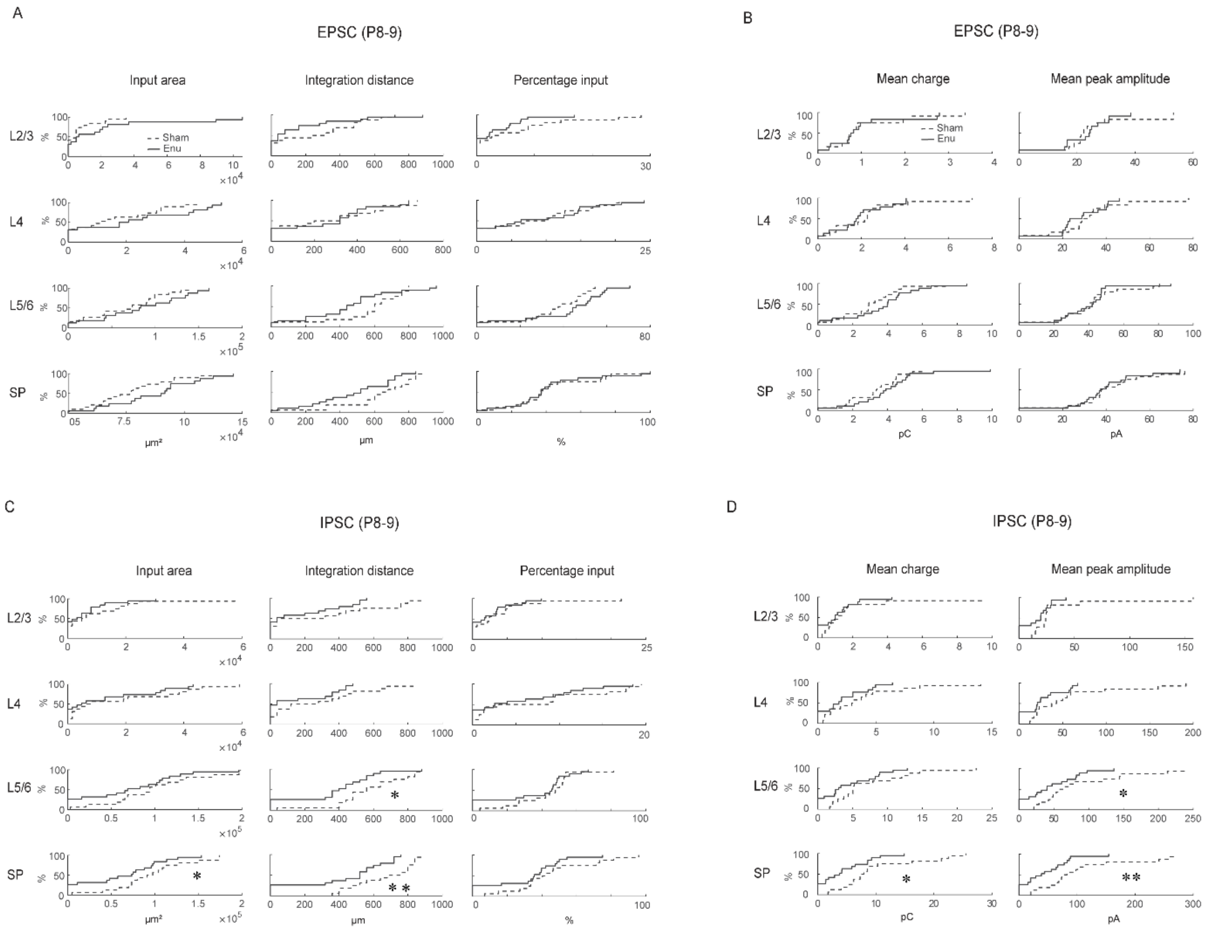


**Figure S3. Laminar measures and connection strengths of excitatory and inhibitory connections after enucleation at P8-9**

**A.** CDFs showing input area, integration distance and percentage input of excitatory connections to SPNs in sham control and enucleated pups at P8-9. **B.** CDFs showing mean charge and peak amplitude of excitatory connections to SPNs in sham control and enucleated pups. **C.** CDFs showing input area, integration distance and percentage input of inhibitory connections to SPNs in sham control and enucleated pups. **D.** CDFs showing mean charge and peak amplitude of inhibitory connections to SPNs in sham control and enucleated pups.

### Figure S4

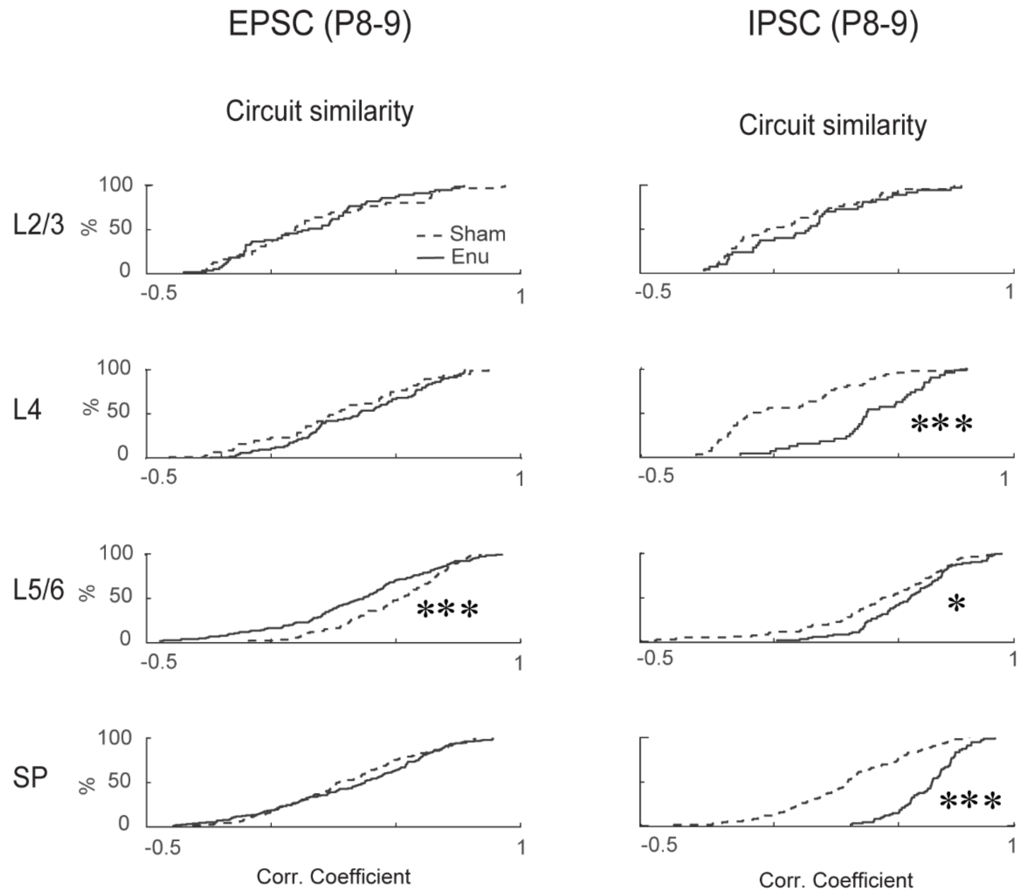


**Figure S4. Connection strength of excitatory and inhibitory inputs after enucleation at P12-15**

**A.** CDFs showing circuit similarity of excitatory and inhibitory connections to the SPNs in sham and enucleated pups at P8-9.

### Figure S5

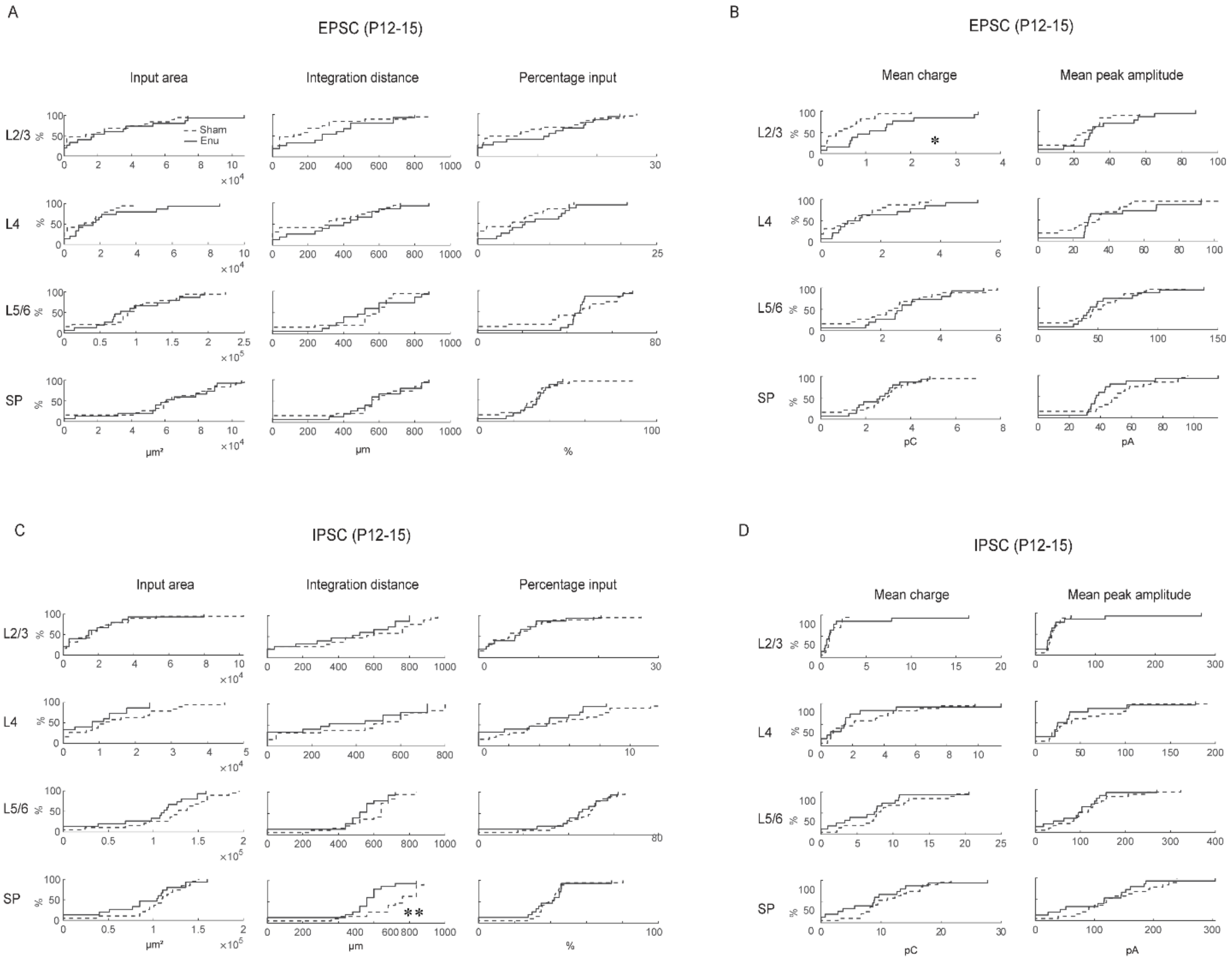


**Figure S5. Laminar measures and connection strengths of excitatory and inhibitory connections after enucleation at P12-15**

**A.** CDFs showing input area, integration distance and percentage input of excitatory connections to SPNs in sham control and enucleated pups at P12-15. **B.** CDFs showing mean charge and peak amplitude of excitatory connections to SPNs in sham control and enucleated pups. **C.** CDFs showing input area, integration distance and percentage input of inhibitory connections to SPNs in sham control and enucleated pups. **D.** CDFs showing mean charge and peak amplitude of inhibitory connections to SPNs in sham control and enucleated pups.

### Figure S6

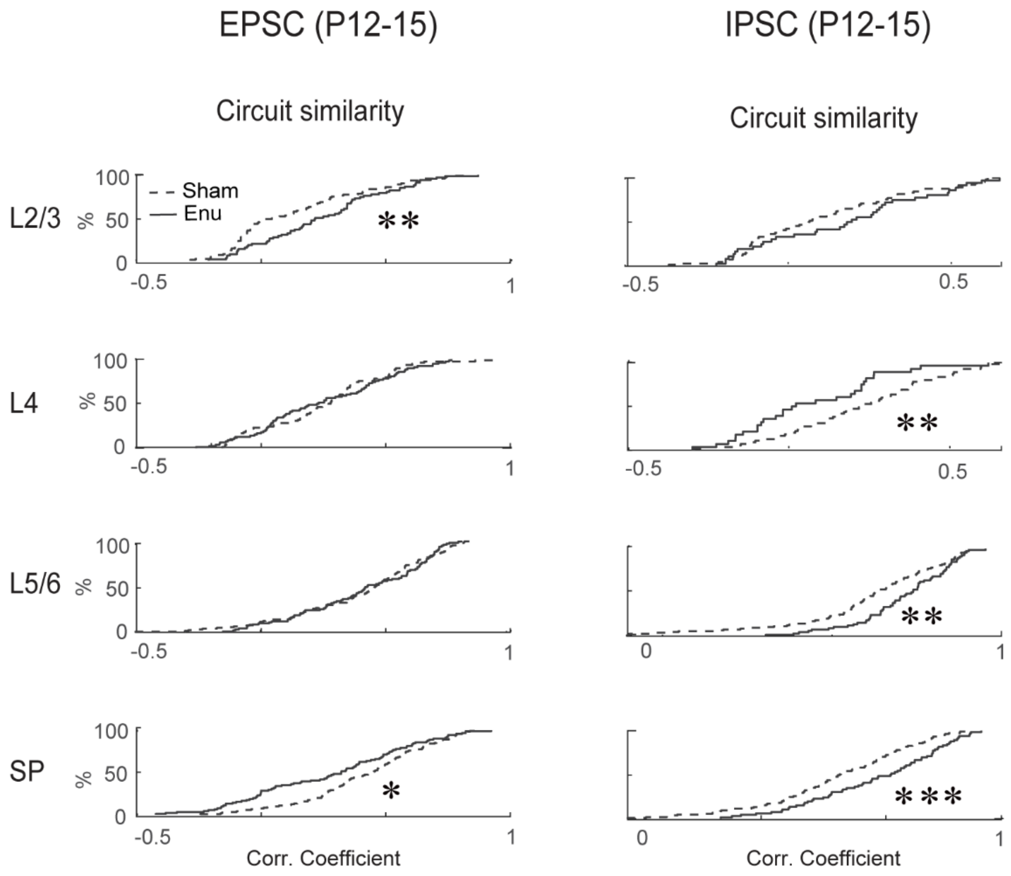


**Figure S6. Connection strength of excitatory and inhibitory inputs after enucleation at P12-15**

**A.** CDFs showing circuit similarity of excitatory and inhibitory connections to the SPNs in sham and enucleated pups at P8-9.
